# Batch Effect Correction to Enhance Utility of a Publicly Available ALS Proteomics Dataset

**DOI:** 10.64898/2026.10.27.762202

**Authors:** Grace Cooper, Inez Oh, Philip Payne, Zachary Abrams, Timothy Miller

## Abstract

**Background:** Large, publicly available databases of patient data have been successful in identifying associations between neurodegenerative diseases and biological features that might otherwise go unnoticed. The Answer ALS Consortium is one such database that has generated a rich repository of multi-omic data from people with amyotrophic lateral sclerosis (ALS). However, large studies can suffer from batch effects that obscure genuine biological signal and prevent experiment reproducibility. In this study, we identified a batch effect in the Answer ALS 1-2 proteomic dataset (508 samples x 3,643 proteins) and corrected the effect using both the internal controls provided by Answer ALS and the batch effect-correction algorithm (BECA) ComBat.

**Results:** We identified a batch effect in the Answer ALS Dataset 1-2 proteomic dataset that, uncorrected, may obscure biology. Proteins MOS and H4 were outliers of expression in the dataset that correlated with the batch effect. Removing MOS and H4 did not correct the effect fully but did reveal new principal component (PC) contributing proteins related to motor neurons and potentially ALS: AINX, NFL, and NFM. The batch differential controls (BDCs) provided by Answer ALS were used to correct samples in their respective batches. ComBat, a well-characterized BECA, was applied to the dataset both before (ComBat-BDC) and after (BDC-ComBat) correction with the BDCs. BDC-ComBat was the only method that successfully reduced batch effect across the entire dataset. This method also preserved four of the top six PC-contributing proteins from the original dataset, including three neuronal proteins, suggesting that the corrected dataset recapitulates biology as well as the variability of the original dataset.

**Conclusion:** Altogether, we produced a version of the Answer ALS proteomic dataset (354 samples by 3,641 proteins) with globally reduced batch effect and protein-level reduced batch effect. This version of the dataset recapitulates the individual protein variability and important neuronal proteins of the original. We also created a workflow for identifying and correcting batch effects that can be applied to other large data repositories.

## Background

Amyotrophic lateral sclerosis (ALS) is a multi-system neurodegenerative disorder characterized by the progressive loss of upper and lower motor neurons in the central nervous system (CNS) (1). Approximately 5,000 people are diagnosed with ALS every year in the United States (2). People with ALS experience muscle weakness, muscle atrophy, and eventual paralysis (2). ALS is nearly universally fatal, and the average survival time between symptom onset and death is three years (3). With the exception of tofersen for *SOD1* genetic ALS, FDA approved drugs have a modest impact on disease (4–6). There is no cure for ALS, and people are primarily treated through supportive care and symptom management (7). The lack of effective treatments for ALS engenders an urgent need to find new potential drug targets. Creating large, publicly available databases of ALS patient samples has become a common approach to increase statistical power for identifying associations between ALS genetics and pathophysiology (8–13). This is a cost-effective, practical way for many research groups to access large amounts of ALS patient data. These databases have been successful; at least nine genes have been associated with ALS in the last ten years using these types of resources (14–22).

One recently created ALS database is the Answer ALS Neuromine Data Portal, a product of the Answer ALS Consortium (8). Individuals were recruited across six different clinical sites, then blood samples and longitudinal clinical data were collected from 1,269 total non-disease control participants and participants with ALS. These blood samples were reprogrammed into induced pluripotent stem cells (iPSCs) and then differentiated into motor neurons. Multi-omic datasets (genomic, epigenomic, transcriptomic, proteomic) were generated from these motor neuron samples. Few studies so far have used Answer ALS, and much discovery is yet to be done on the aggregated datasets (23–27).

Despite their many benefits, large datasets collected over time like Answer ALS are vulnerable to batch effects (28–31). Minor variations in protocols for sample preparation, reagents, sequencing instruments, sites for sequencing, and computational processing platforms can all contribute to batch effects within a dataset (32). The multi-step differentiation process Answer ALS used to generate the multi-omic datasets makes these data inherently vulnerable to batch effects. Batch effects are problematic because they can lead to false correlations, obscure genuine biological signals, and prevent successful reproduction of an experiment (33,34). Further, these effects can extend to any type of omics data, including proteomics (33–35).

Answer ALS has already been informed that a batch effect exists in their proteomic dataset, and samples have been removed in accordance with the discovery (36). It has yet to be determined if there is a batch effect in the second iteration of the proteomic dataset (Dataset 1-2). Therefore, we aimed to determine if a batch effect exists in the Answer ALS 1-2 proteomic dataset and, if found, correct this effect.

The Answer ALS Consortium currently provides metadata for two steps of their data generation process: the differentiation of iPSCs into motor neurons, termed ‘batches’, and the omic sequencing of said motor neurons, termed ‘shipments’ (36). We refer to the differentiation batches as ‘iPSC Motor neuron Differentiation Batches’ (iMDBs) to avoid confusion with the general term ‘batch effect’. All iMDBs and shipments contain a biological replicate of a non-disease control sample to control intra-batch variability (8). We used these biological replicates to adjust levels of protein expressions for samples within each iMDB. Because this method was not sufficient to reduce batch effect, the well-validated batch effect-correction algorithm (BECA) ‘ComBat’ (37,38) was also applied to the dataset. These two methods of batch effect correction were applied to the Answer ALS proteomic dataset separately and sequentially.

## Methods

### Data Generation and Acquisition

Data used in this study were obtained from the Neuromine Answer ALS Data Portal (AALS-01184). Proteomic analysis was performed on the ‘pre-batch corrected protein matrix’, which contains 3,643 proteins as features. This is referred to as ‘Level 4 data’ in the Answer ALS data portal. Feature reduction and other pre-processing steps were performed as previously described prior to data download (8). Proteomes and corresponding clinical data were downloaded from the Answer ALS data portal from the March 2024 data release (36). Additional control samples were obtained through direct contact with the program director of Answer ALS, for a total of 508 samples. The proteomes for these samples were generated through the process outlined in **Supplemental Figure 1**. ‘iMDBs’ and ‘shipments’ are two different, yet related, metadata variables for groups of samples formed in two steps of the data generation process. We chose to focus on iMDBs because the differentiation of iPSCs to motor neurons occurred prior to sequencing. Therefore, any batch effects that impact the iMDBs would impact shipments as well. **Supplemental Table 1** shows the ‘Sample Mapping File February 2024’ available from Answer ALS, which contains the complete list of samples in iMDBs and shipments (36). Additionally, a biological replicate of one non-disease female control, ‘NEUEU392AE8’, was differentiated with each iMDB and sequenced with each shipment to ensure consistency between groups (8). These are referred to as batch differential controls (BDCs) for the iPSC-to-motor neuron differentiation.

### Dataset Filtering

Clinical data and metadata associated with the proteomic dataset were used to filter samples outside of the experiment parameters (**Supplemental Figure 2**). In terms of feature-based filtering, the number of proteins that were not detected in each proteomic sample was calculated. If 33.33% or more of the total proteins were not detected in a sample, the sample was excluded. The remaining samples were log-transformed and visualized via boxplot in two rounds; outliers were also excluded. Specific details about sample exclusions can be found in **Supplemental Table 2**. After all filters were applied, 430 proteomic samples remained in the final dataset that was used for all further analyses.

### Characterizing iMDBs

The filtered proteomic dataset contained 305 ALS samples and 125 non-disease control samples. **Figure 1** shows the distribution of these samples across iMDBs and sequencing shipments. iMDB numbers start at 10 and shipment numbers start at 4 due to a previously identified batch effect in iMDBs 1-9, which were subsequently removed from the Answer ALS data portal (36). We removed iMDBs 46 and 49 entirely for globally low protein values and discordant samples, respectively (**Supplemental Table 2**). All information in the remainder of this paragraph was taken from personal communication with the Program Director for OnPoint Scientific, Dr. Terri Thompson, between the dates of March 30^th^ and April 3^rd^, 2026. iMDBs 53 and 56 were run as a replication cohort and proteomics was not included in that analysis. Shipment 19 and iMDB 14 had low or nonexistent sample numbers because they were processed by a separate facility. iMDB 57, on the other hand, was not in the Sample Mapping File for any omic data type because that iMDB was the second derivation of the shipment level technical controls. Shipment 18 encompassed only iMDB 55 while other shipments encompassed at least three iMDBs because of supply chain issues. Shipments 13 and 14 were comprised of different sample distributions despite being generated close together temporally. This was due to the inclusion of external controls for Shipment 14 and subsequent iMDBs/shipments. The low number of consenting controls in the original study necessitated the inclusion of ∼200 outside control samples.

**Figure 1:**
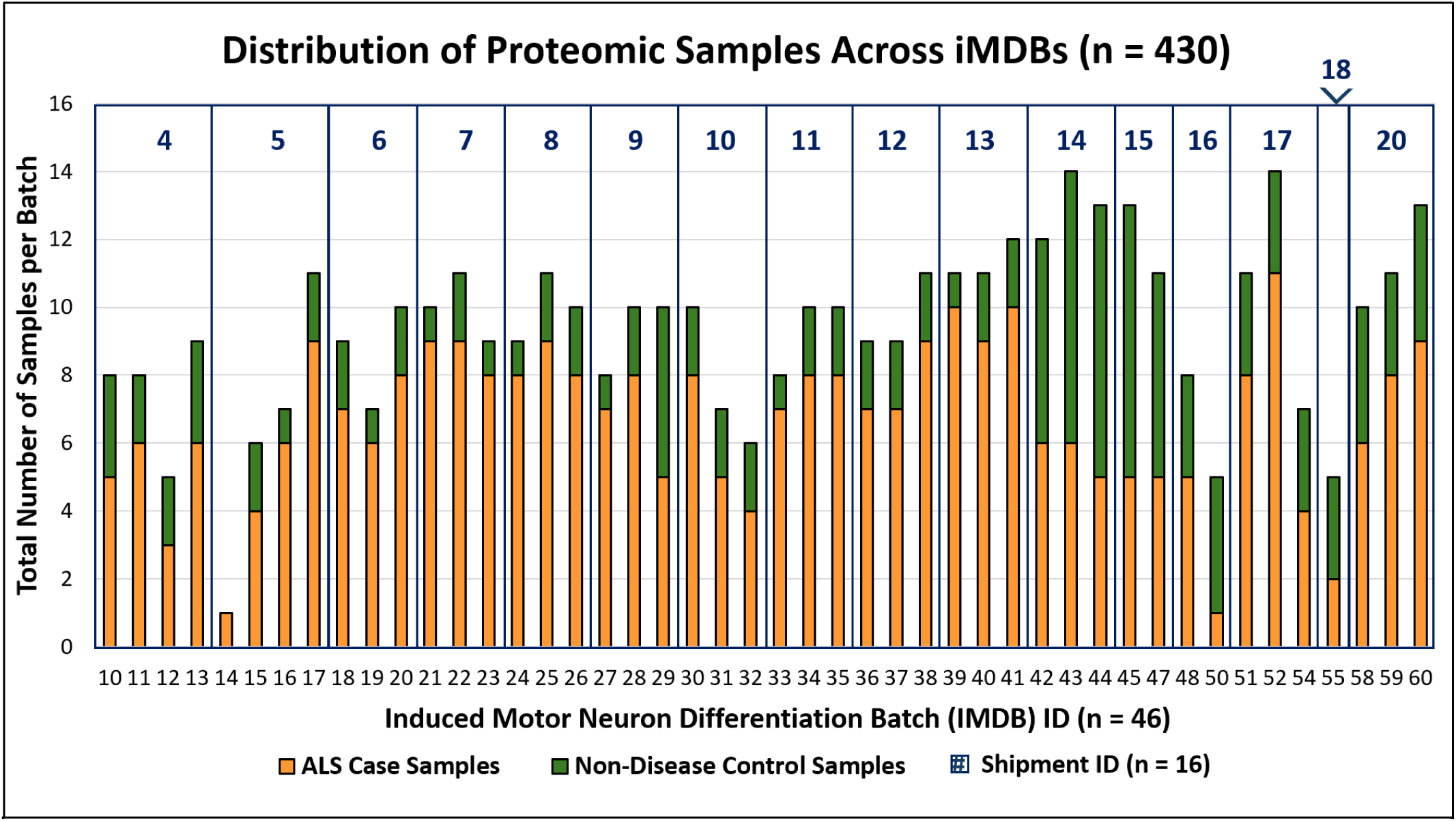
Proteomic samples are unevenly distributed across iMDBs and shipments. Proteomic samples were plotted by iMDB, sequence processing shipment, and sample type (case or control).

### Principal Component Analysis (PCA) Generation

R version 4.4.3 “Trophy Case” (2025-02-28) was used for all analyses. Principal components were generated using the ‘prcomp’ function from the base R package ‘stats’. Creation of PCA plots was performed with R packages ‘ggplot2’ (39) and ‘ggfortify’ (40). Shipment ID was chosen as a metadata factor proxy for iMDB ID, for ease of interpretation in the PCA plots. 16 color-coded shipments are easier to discern than 47 color-coded iMDBs. For the PCAs that visualize protein expression, quartile values 1-4 for the protein were determined from all samples in the dataset and used to color-code these samples.

### Determination of PC-Contributing Features

The contribution of proteins to PC1 and PC2 for a given analysis were determined by examining the output of function ‘prcomp’. The function of the proteins and corresponding gene names were annotated with the UniProt Protein Knowledgebase and GeneCards: the Human Gene Database (41,42). The relationship of each protein to ALS disease pathology was explored using the Amyotrophic Lateral Sclerosis online Database (ALSoD) (43), the Adult Genotype-Tissue Expression (GTEx) project (44), the Reactome Pathway Knowledgebase (45), and Uniprot.

### Individual Protein Testing

We examined all proteins in different iterations of the proteomic dataset to determine if the observed effects extended to individual proteins. The Pearson correlation coefficient ‘*r*’ (46) between iMDB ID and each protein’s value per sample was calculated for each protein. The null hypothesis was that iMDB ID and protein values are unrelated and should generate a small ‘*r’*. Therefore, we categorized a protein with a Pearson correlation coefficient (PCC) above 0.25 or below -0.25 as ‘impacted by batch effect’. This allowed us to determine the number of proteins in each dataset impacted by batch effect. This analysis was performed for every iteration of the dataset, as shown in **Table 1**. We also randomly selected 10 features from the examined subgroup and visualized them for every dataset (**Supplemental Figures 3-12**).

**Table 1:** The number of proteins impacted by batch effect in each iteration of the proteomic dataset. Proteins were determined to be impacted by batch effect if the Pearson correlation coefficient (PCC) between protein values and iMDBs was > 0.25 or < -0.25.

| **Dataset (**samples x proteins**)** | **Number of Proteins Impacted by Batch Effect** |
|---|---|
| **Raw proteomic dataset** (430 x 3,643) | 1,971 (54.1%) |
| **BDC-corrected dataset** (380 x 3,641) | 1,487 (40.8%) |
| **BDC-ComBat-corrected dataset** (354 x 3,641) | 537 (14.7%) |
| **ComBat-BDC-corrected dataset** (293 x 3,641) | 1,258 (34.6%) |

### The ComBat Algorithm

ComBat is a batch effect-correction algorithm (BECA) from the ‘sva’ R package with three basic steps (37,38). First, the dataset is standardized by individual features so that features have overall similar means and variances. Empirical Bayes batch effect parameters are then estimated using either parametric or non-parametric empirical prior estimates. Then, the calculated adjusted batch effect parameters are used to adjust the standardized dataset. ComBat correction was performed nonparametrically without covariates in R version 4.4.3 “Trophy Case” (2025-02-28) for all analyses.

## Results

### Confirmation of Batch Effect

To determine if batch correction is necessary before applying a correction technique, we averaged the expression of every protein (n = 3,643) together for each sample and then visualized samples in their respective iMDBs (**Figure 2A**). A batch effect was identified, as individual iMDBs were not centered around the overall protein expression average (n = 74,535.6). In some iMDBs, all samples showed above average levels of protein expression, while in other iMDBs, all samples were below average. A PCA plot was also generated using this dataset and subsequently color-coded by sample shipment ID (**Figure 2B**). Samples were grouped in this PCA plot by shipment/iMDB. Since shipments contain 2-3 iMDBs, a batch effect that affects shipments affects iMDBs as well. PC1 and 2 also account for 91.17% of the total variance within the dataset. Although the average expression is useful for viewing the shape of the dataset, using only averages can obscure effects on the individual feature level. We determined that 1,971 (54.1%) individual proteins were impacted by this batch effect (**Table 1**). Individual protein visualizations are available in **Supplemental Figures 3-12**. These results together show that there is a batch effect in the Answer ALS 1-2 proteomic dataset and that the effect is both global and individual.

**Figure 2:**
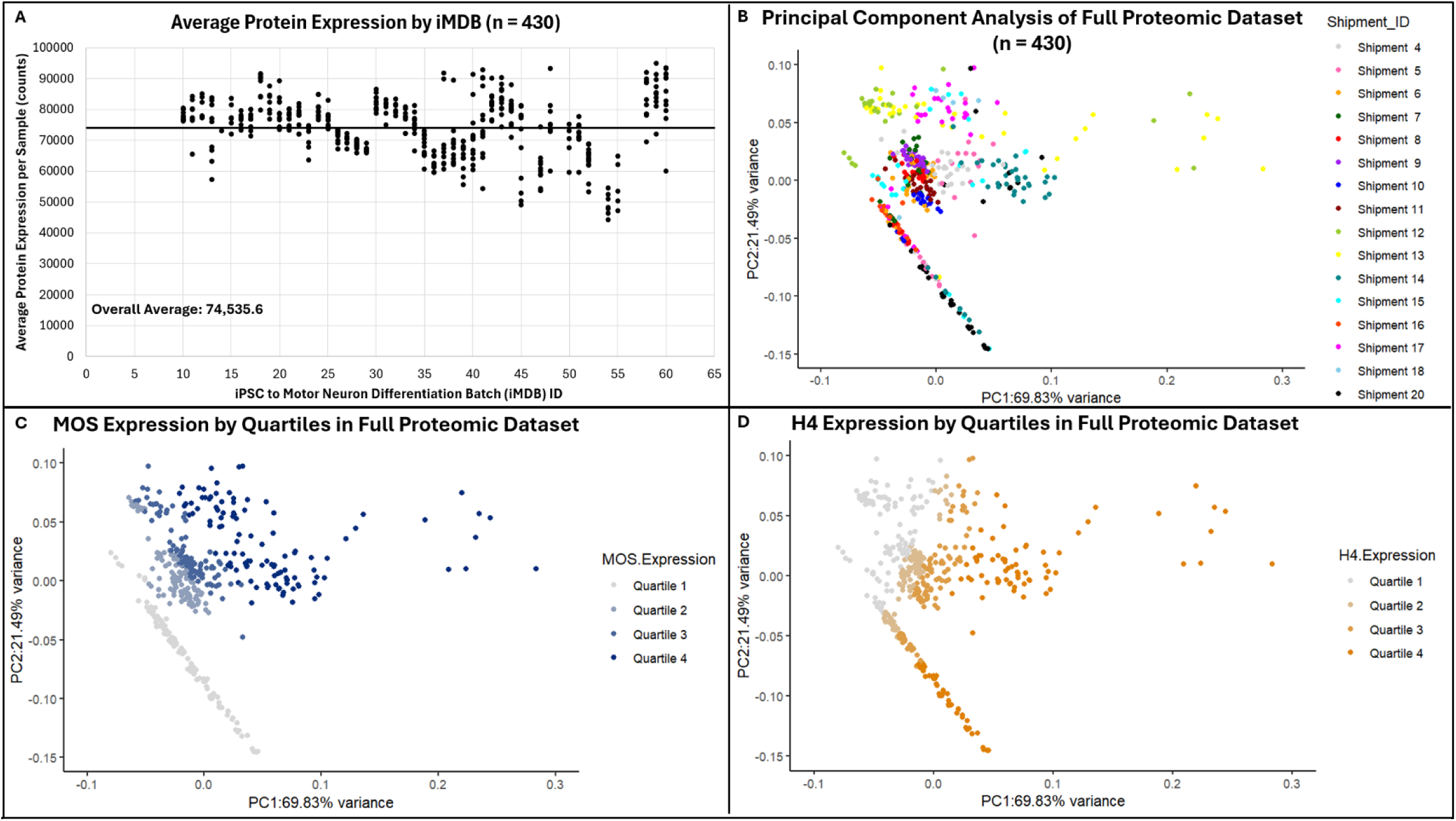
Expression of proteins ‘MOS’ and ‘H4’ correlate with batch effect in the raw proteomic dataset. A) Average protein expression per sample plotted by iMDB ID. B) PCA color-coded by shipment ID. C) PCA color-coded by MOS protein expression, divided into quartiles. D) PCA color-coded by H4 protein expression, divided into quartiles.

Additionally, we found that proteins ‘MOS’ and ‘H4’ were major contributors to PC1 and 2. MOS is a serine/threonine kinase involved in MAPK signaling, and H4 is Histone 4, a core component of the nucleosome (41). Neither protein is known to be related to ALS pathology, and neither of them are highly expressed in the central nervous system (41,43,44). MOS contributed most highly to PC2 (0.782) and quartile 1 expression of MOS (no measured expression) also correlated with the linear structure in this PCA plot. H4 contributed most highly to PC1 (0.796). When plotted, MOS (**Figure 2C**) and H4 (**Figure 2D)** expression levels correlated highly with both PCs. MOS and H4 expression levels were also visualized across the iMDBs in the original dataset and the variation of these proteins suggests the presence of batch effects (**Supplemental Figure 13**). Together, these analyses suggest that MOS and H4 expressions are correlated to batch effect within the proteomic dataset.

### Removal of MOS and H4

To determine whether MOS and H4 expression levels were correlated with or causing the batch effect, we removed MOS and H4 from the dataset completely (n = 3,641) and re-ran the proteomic dataset analysis (**Figure 3**). We found that the removal of these two features dropped the overall average protein expression per sample from 74,535.6 to 66,200.8; however, the shape of the dataset did not change (**Figure 3A**). In contrast, the PCA plot of the dataset without MOS and H4 showed a looser spatial arrangement of samples than the original PCA plot and did not contain the linear structure formed by MOS (**Figure 3B**). Shipments/iMDBs still grouped together within the looser spatial arrangement of the PCA plot. This suggests that while MOS and H4 expression levels correlate and/or contribute to the batch effect, they are not the sole cause.

**Figure 3:**
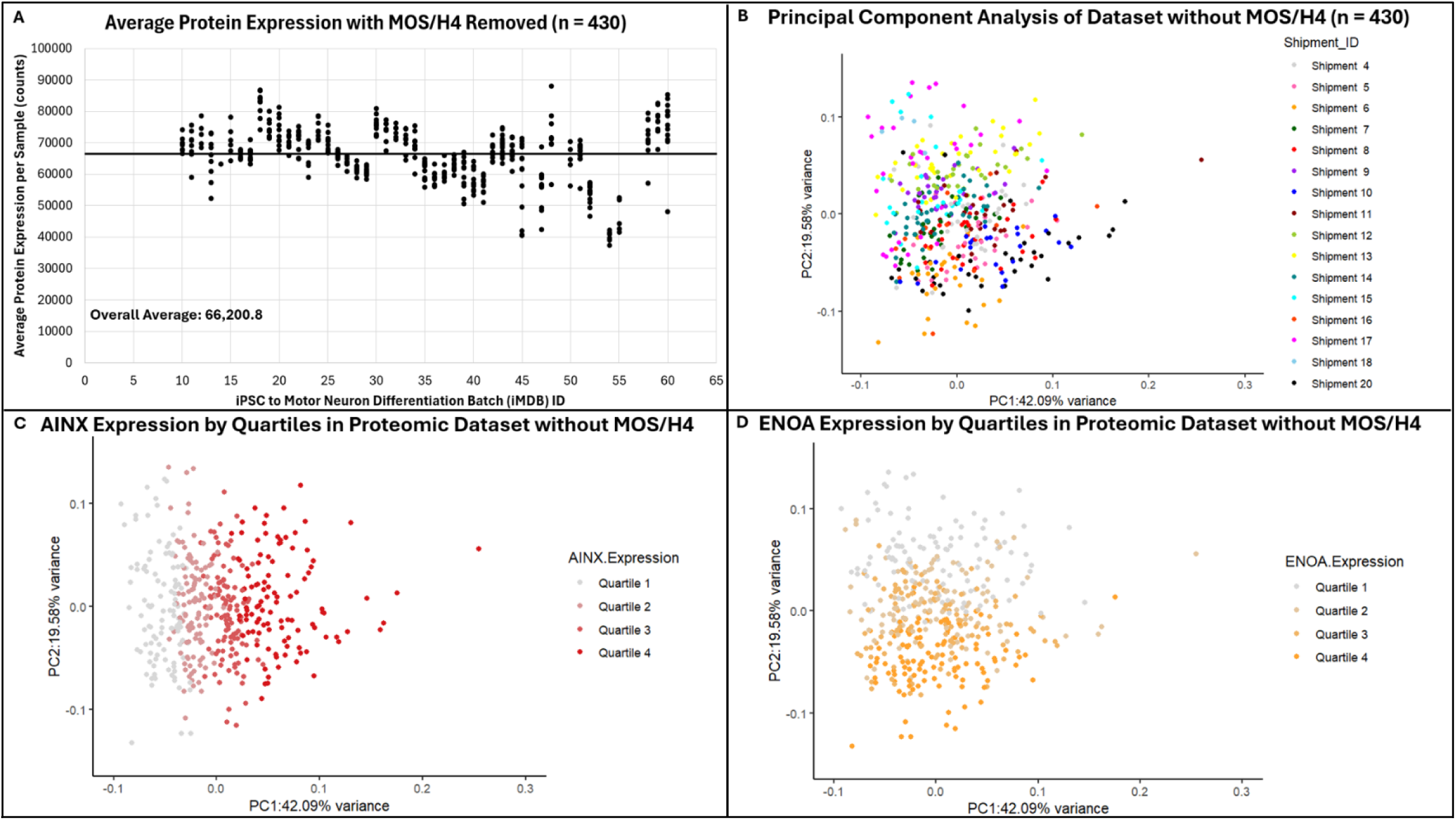
Removal of MOS and H4 reveals PC-contributing neuronal proteins but does not remove batch effect. All experiments performed in the dataset without MOS and H4. A) Average protein expression per sample plotted by iMDB ID. B) PCA color-coded by shipment ID. C) PCA color-coded by NFL protein expression, the second-most contributing protein in PC1, divided into quartiles. D) PCA color-coded by KCRB protein expression, the second-most contributing protein in PC2, divided into quartiles.

Removing MOS and H4 also revealed a top PC-contributing protein that is neuronal. The second-highest contributor to PC1 was protein AINX (**Figure 3C**), which is involved in the maintenance of neurons and highly expressed in nervoustissues (41,44). The second highest contributor to PC2 is protein ENOA (**Figure 3D**). This protein is not known to be neuronal and is not highly expressed in nervous tissues (41,44). Nonetheless, the high contribution of the neuronal protein AINX to PC1 suggests that the removal of MOS and H4 from the dataset reveals genuine biology. We therefore chose to exclude proteins MOS and H4 from all further analyses.

### BDC Batch Correction

With MOS and H4 removed from the dataset, we next wanted to correct for batch effect across the whole dataset using the BDCs provided by Answer ALS. However, we identified inconsistencies with the current BDCs that impacted the batch correction process. iMDBs 13, 23, 39, and 59 did not have BDCs, and iMDB 14 contained only a single sample. Additionally, there was no BDC for iMDB 54 in the Sample Mapping File (**Supplemental Table 1**). We removed the iMDBs without BDCs, which reduced the dataset from 430 samples to 382 samples. We then visualized the average protein expression for BDCs (pink) and other samples across this smaller subset of the dataset (**Figure 4A**). We chose to move forward with BDC correction despite these difficulties because often the most successful correction methods are biologically relevant to the original dataset and minimally transformative, and BDC correction fits both criteria (47).

**Figure 4:**
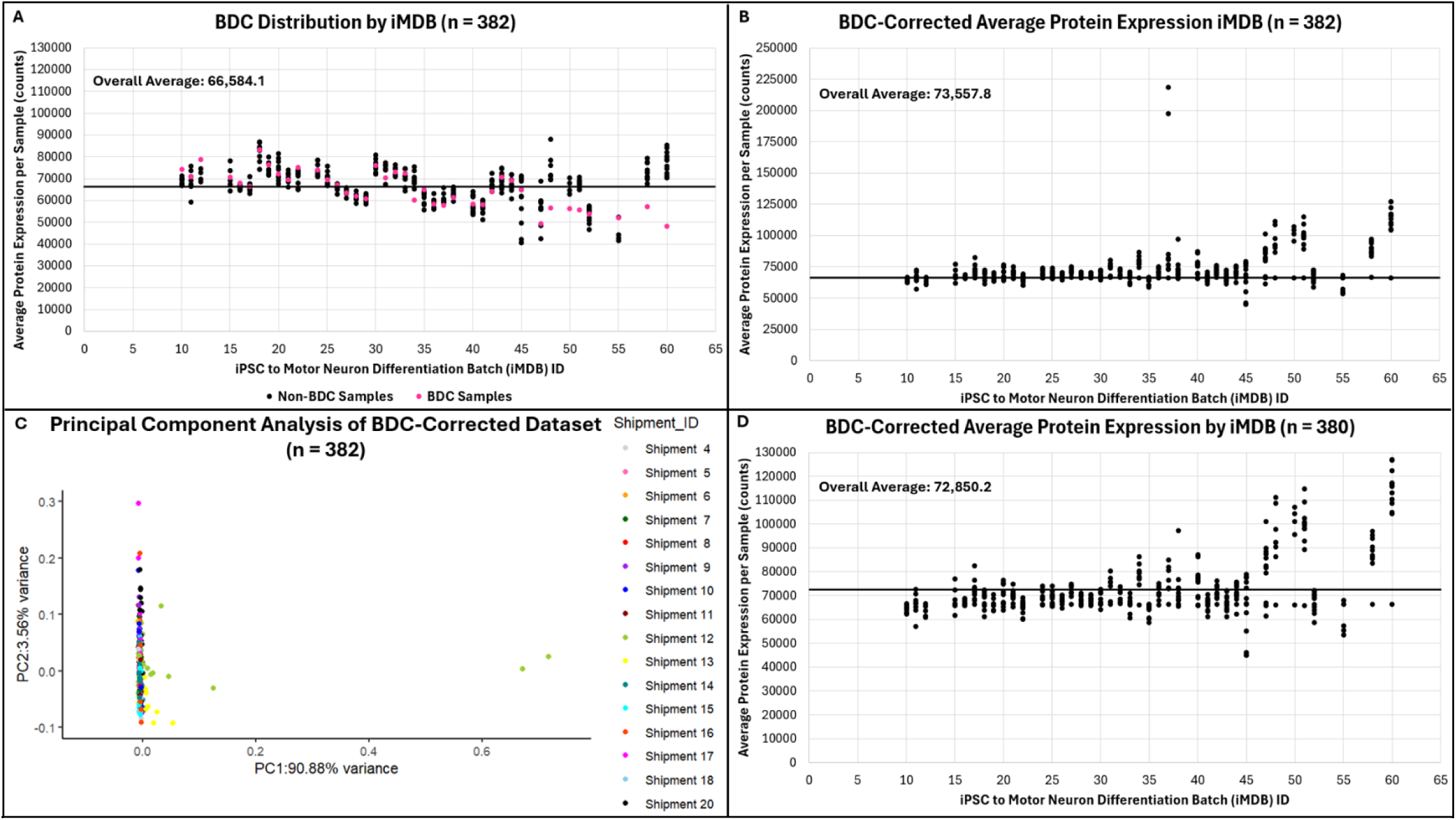
BDC correction reveals sample outliers and unevenly impacts the later iMDBs. (**47–60**). A) Average protein expression per sample color-coded to show BDCs (pink) and all other samples (black). B) Average protein expression after correcting iMDBs with BDCs. C) PCA of BDC-corrected dataset color-coded for shipment ID. D) BDC-corrected average protein expression plot with sample outliers removed.

To correct batch effect with the BDCs, we first took the average of every individual protein (n = 3,641) across all samples (n = 382). We then divided the average by the BDC value for the given protein in each iMDB. This produced a ‘factor’, a number greater than 0, that represents how much the BDC was adjusted to reach the average expression of that protein. Then, we multiplied every other sample within that particular iMDB by the BDC-produced ‘factor’. This process was repeated for every protein, in every iMDB. **Figure 4B** shows the BDC-corrected average protein expression per sample across all iMDBs. The overall average increased from 66,200.8 to 73,557.8 after BDC correction. The correction also produced two outlier samples, both in iMDB 37 (**Figure 4B**). Case ‘NEUAE228FF6’ (FF6) is a male ALS case sample with an average protein expression 13.79 times greater than the standard deviation (15,838.94) and 2.97 times greater than the overall average protein expression. Control ‘NEUNC876ZB2’ (ZB2) is a male non-disease control sample with an average protein expression 12.47 times greater than the standard deviation and 2.69 times greater than the overall average protein expression. These outliers were also identified in the PCA plot of the BDC-corrected dataset (**Figure 4C**). Although both samples are in iMDB 37, the other samples in this iMDB did not qualify as outliers, so only ‘FF6’ and ‘ZB2’ were removed from the dataset. **Figure 4D** shows the BDC-corrected average protein expression in the remaining 380 samples. The overall average protein expression dropped from 73,557.8 to 72,850.2. We determined that 1,487 (40.8%) individual proteins in this dataset were impacted by batch effect (**Table 1**). Individual protein visualizations are available in **Supplemental Figures 3-12**.

Once the outliers were removed, the difference in average protein expression between iMDBs 10-45 and 47-60 became apparent (**Figure 4D**). Samples in the corrected iMDBs 10-45 had average protein expressions mostly between 60,000 and 80,000. Samples in the corrected iMDBs 47-60, on the other hand, had average protein expressions mostly above the average (72,850.2), and as high as 126,911.4, which is 1.74 times greater than the average. We attribute this difference to the variability in the uncorrected BDC average protein expressions. Prior to correction, BDCs in iMDBs 10-45 matched the rest of the samples in the iMDBs; when the iMDB was above the overall expression average, so was the BDC (**Figure 4A**). After iMDB 47, the average protein expression of the BDCs were all below the overall expression average regardless of the rest of the samples in their respective iMDBs. Correcting with these BDCs resulted in artificially inflated average protein expressions in their respective iMDBs and exacerbated the batch effect between the first and last iMDBs.

Another sample ‘NEUWD946RDA’ was identified as an outlier and removed in subsequent analyses (**Supplemental Figure 14**), bringing the number of remaining samples to 379. Attempts to reduce batch effect by removing top PC-contributing proteins (**Supplemental Figure 15**) and outlier iMDBs (**Supplemental Figure 16**) alone or in combination (**Supplemental Figure 18**) were not successful. Altogether, these results show that correcting with the built-in BDC samples is not sufficient for reducing batch effect across the entire dataset.

### ComBat Selection Rationale

Our lack of success using the BDCs to reduce batch effect across the proteomic dataset necessitated an additional correction method. We have already established that the proteomic dataset has small batch sizes (<15 samples) and an uneven distribution of sample types across iMDBs, two characteristics that limit the batch correction methods that can be applied to the datasets (33,37,48). ComBat is well-suited for handling these concerns and is extensively used for correcting batch effect (37,38). Additionally, ComBat is easy to implement and runs quickly compared to other batch effect-correction algorithms (BECAs) (49). We hypothesized that applying ComBat in combination with BDC correction to the proteomic dataset would preserve the correction of the earlier iMDBs (10–45) and adjust the artificially inflated samples in the later iMDBs (47–60).

### Combined Batch Corrections

ComBat correction was performed on the previously generated BDC-corrected dataset (379 samples x 3,641 proteins). **Figure 5A** shows the average protein expression across the iMDBs in the BDC-ComBat-corrected dataset. The overall average protein expression per sample was 72,804.7, close to the overall average for the BDC-corrected dataset (72,850.2). While the overall average barely changed, the shape of the BDC-ComBat-corrected dataset was less impacted by batch effect. A PCA plot of this dataset showed that iMDBs 45 and 47 (shipment 15) were outliers in PC2 (**Figure 5B**). 10 of 24 samples (41.7%) in these batches were visibly separated from the rest of the dataset. These iMDBs/shipment were removed entirely due to the high percentage of outlier samples and the principal component analysis was re-run without them (**Figure 6A**), creating a dataset with 354 proteomic samples. The resulting PCA plot was the first produced through the batch correction process where samples were no longer visibly grouped by iMDB/shipment. These two figures together show that correcting with BDCs and then ComBat successfully reduces batch effect across the entire dataset.

**Figure 5:**
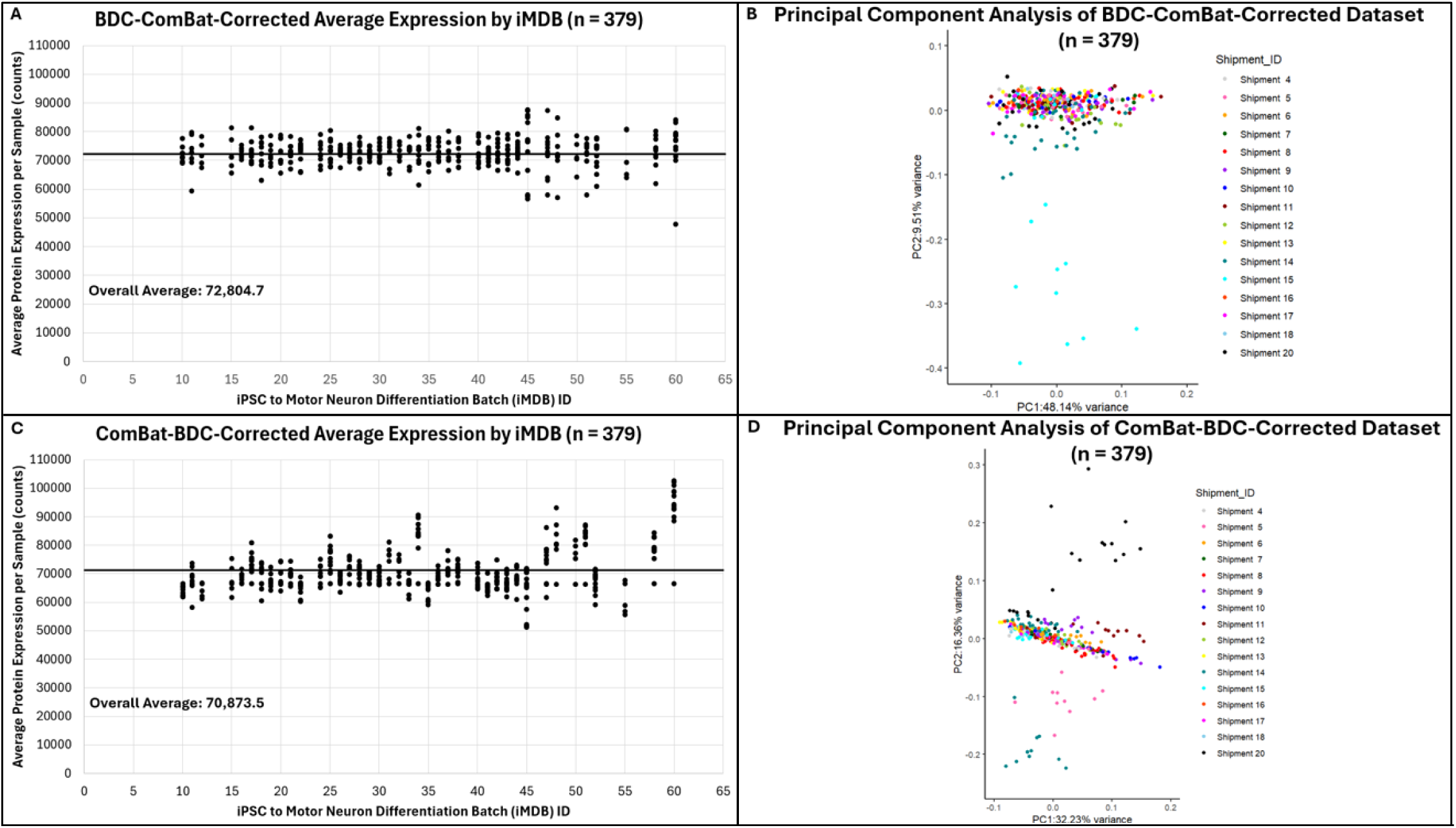
Combining ComBat and BDC correction reduces batch effect if BDC correction is done first. BDC-ComBat-corrected A). Average protein expression by iMDB ID and B). PCA plot color-coded by shipment ID, ComBat-BDC-corrected C). Average protein expression by iMDB ID and D). PCA plot color-coded by shipment ID.

**Figure 6:**
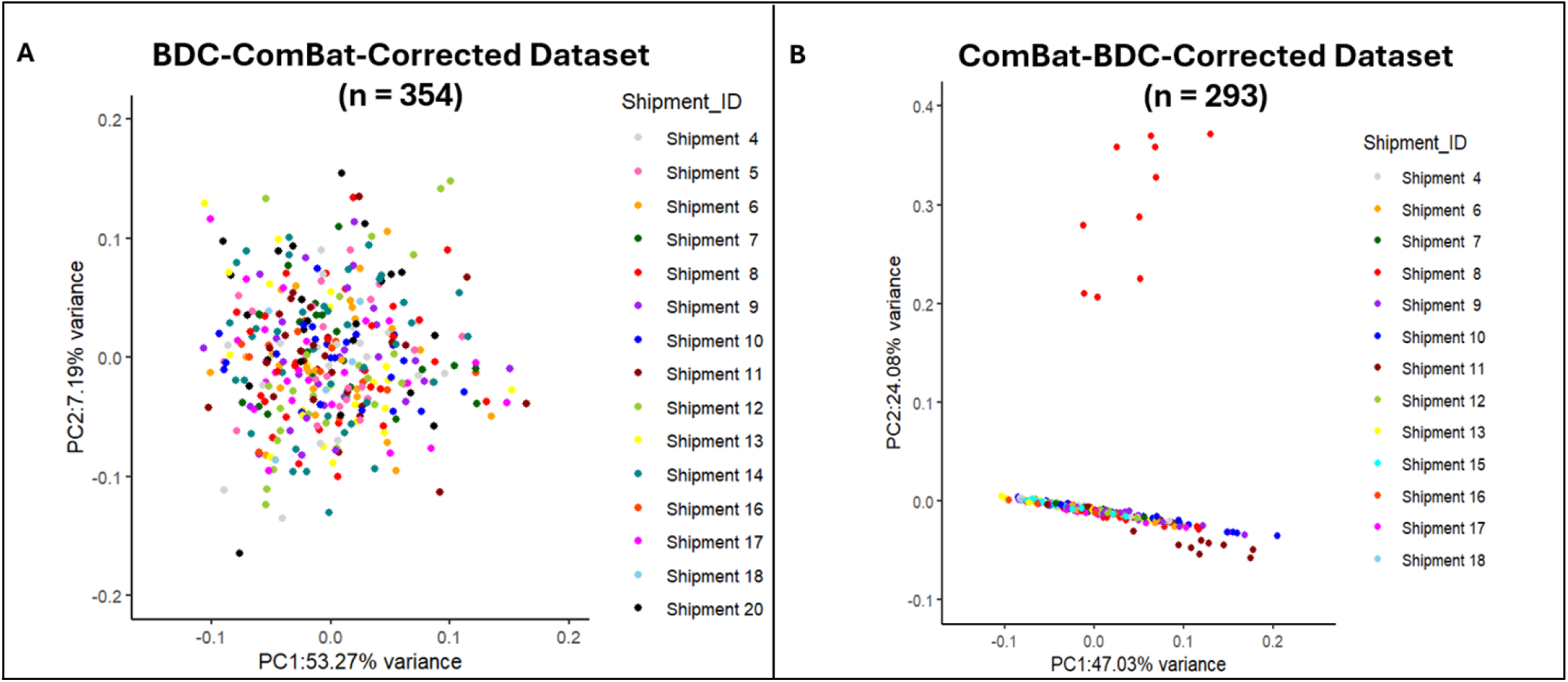
Correcting with BDCs and then ComBat reduces batch effect while preserving more of the dataset than the reverse. A) PCA plot of the BDC-ComBat-corrected dataset color-coded by shipment ID. B) PCA plot of the ComBat-BDC-corrected dataset color-coded by shipment ID.

With these results, we wanted to investigate if the sequential order of BDC and ComBat correction was integral to reducing batch effect. A dataset was produced with matched samples to the BDC-corrected dataset (n = 379), but with raw proteomic values. This allowed direct comparison to the BDC-ComBat-corrected dataset without the confounding factor of different sample numbers. ComBat correction was performed on this dataset, and then the dataset was corrected with BDCs as described above. **Figure 5C** shows the average protein expression across the iMDBs in the ComBat-BDC-corrected dataset. The overall average protein expression per sample was 70,873.5. Unlike the BDC-ComBat-corrected dataset, the shape of this dataset recapitulated the issues observed in the BDC-corrected dataset with the later iMDBs (47–60). When a PCA plot was generated from this dataset, it also recapitulated the shape of the BDC-corrected only PCA plot (**Figure 5D**). iMDBs 15-17 (shipment 5), iMDBs 42-44 (shipment 14), and iMDBs 58 and 60 (shipment 20) all contained samples identified as outliers in PC1. 10 of 24 samples in shipment 5 (41.7%), 10 of 39 samples in shipment 14 (25.6%) and 12 of 23 samples in shipment 20 (52.2%) were separate from rest of the dataset. As a result of the high percentage of outlier samples, all three groups of iMDBs/shipments were removed from this iteration of the dataset, and principal component analysis was performed again on the now 293 sample dataset (**Figure 6B**). The batch effect was not reduced in this version of the dataset. Shipment 8 still separated from the others, and the rest of the samples were still grouped together by shipment. This iteration of the dataset also required the removal of 61 more samples than the BDC-ComBat-corrected iteration. These results show that correcting first with BDCs and then with the ComBat function generates a dataset with reduced batch effect and more samples preserved than correcting with ComBat first. Additionally, these results suggest that correcting with BDCs and then ComBat is more effective at reducing batch effect than correcting with just the BDCs.

### Protein-Level Batch Correction

The protein-level batch correction results mirror the global results for both combinations of BDC/ComBat corrections. For the BDC-ComBat-corrected dataset, we determined that 537 (14.7%) proteins were impacted by batch effect, a 2.8-fold reduction from the BDC-corrected dataset and a 3.7-fold reduction from the raw dataset (**Table 1**). The number of proteins impacted by batch effect in the ComBat-BDC-corrected dataset, however, was 1,258 (34.6%), only a 1.6-fold reduction from the raw dataset (**Table 1**). BDC-ComBat correction reduces batch effect on twice as many proteins as ComBat-BDC correction.

We also visualized superoxide dismutase 1 (SOD1) protein expression in the different datasets to show the impact of the combined correction approaches on an established ALS-related protein (50–52) (**Figure 7**). In the original dataset, SOD1 expression was lower in iMDBs 20-30 than the surrounding iMDBs (**Figure 7A**). BDC correction alone greatly inflated SOD1 expressions in iMDBs 47-60 and failed to correct the lower SOD1 expressions in iMDBs 20-30 (**Figure 7B**). Both combined approaches adjusted the low expression in these iMDBs, and both approaches produced nearly identical overall average SOD1 expression values (**Figures 7C and 7D**). However, the BDC-ComBat approach was more effective at reducing the batch effect across all iMDBs (**Figure 7C**). Most samples in iMDBs 20-30 were still below the overall average SOD1 expression in the ComBat-BDC approach (**Figure 7D**). For further evidence, 10 additional and randomly selected proteins were visualized for both datasets and are available in **Supplemental Figures 3-12**. These results, combined with the impacted protein estimates, indicate that the BDC-ComBat approach reduces batch effect on individual proteins more than any other tested approach in this publication.

**Figure 7:**
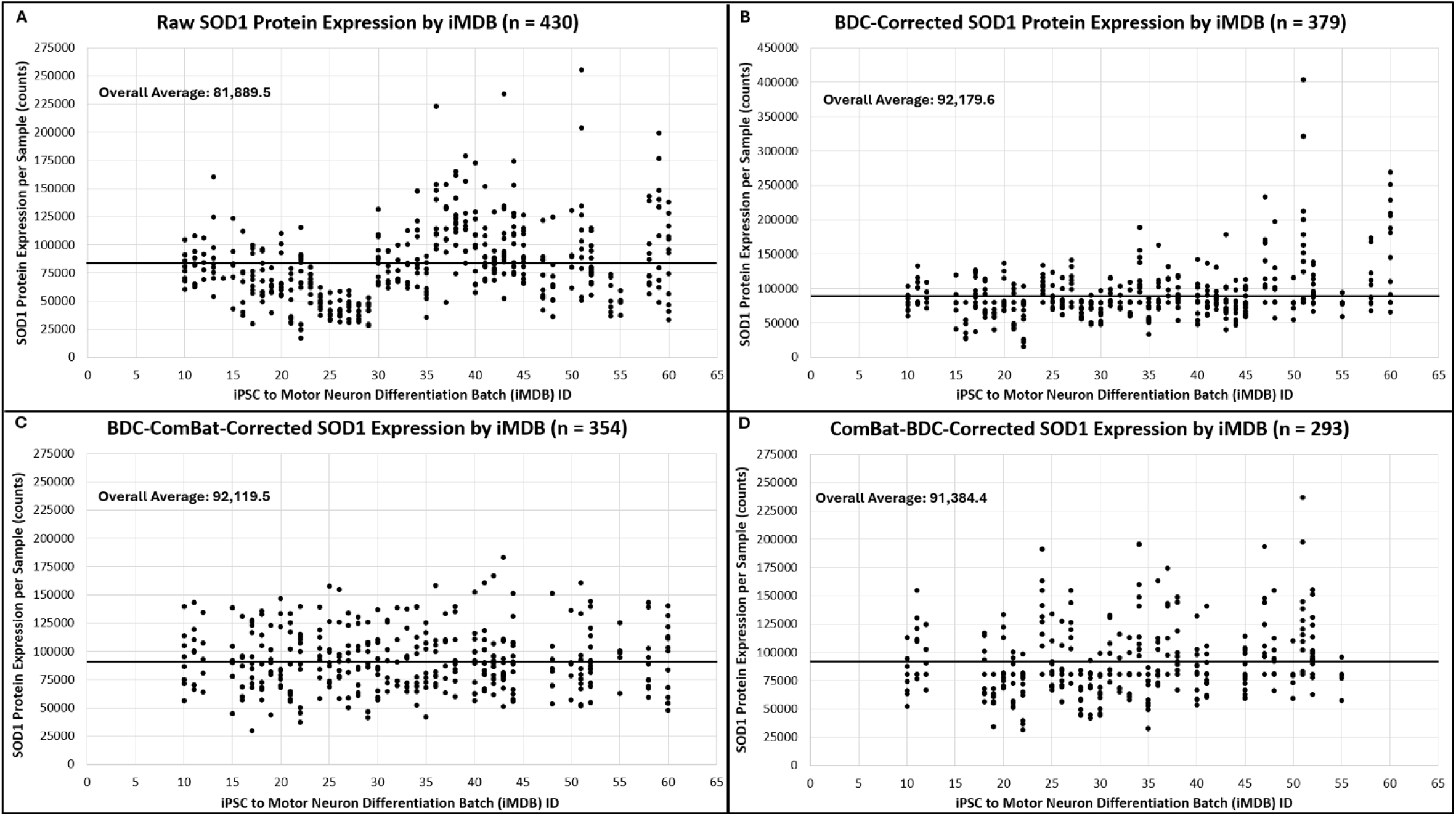
The BDC-ComBat approach corrects for batch effect the best in an ALS-related protein. SOD1 protein expression across all samples is visualized in the A) uncorrected proteomic dataset, B) BDC-corrected dataset, C) BDC-ComBat-corrected dataset, and C) ComBat-BDC-corrected dataset.

### Top-Contributing Proteins

The top three highest-contributing proteins to PCs 1 and 2 were investigated for the BDC-ComBat-corrected dataset to determine if this correction method maintained the biological relevance observed in the uncorrected dataset with MOS and H4 expression removed (**Table 2**). The top three highest-contributing proteins to PC1 were identical in the BDC-Combat-corrected dataset and the uncorrected dataset: AINX (alpha-internexin), NFL (neurofilament light polypeptide), and NFM (neurofilament medium polypeptide). All of these proteins are intermediate filament proteins involved in the morphogenesis and maintenance of neurons (41,44). All are highly and specifically expressed in brain tissue (44). In ALS, the aggregation of intermediate filaments in motor neurons is a pathological hallmark (53). NFL is a biomarker of neurodegeneration (54–57) that, while not specific to ALS, can successfully differentiate ALS from other neurodegenerative diseases in some studies (58–62). Levels of NFL in biofluids have also been found to be higher in ALS than most other neurodegenerative disorders (63,64). NFM is understudied in comparison to NFL in ALS, but there is some evidence that NFM also marks neurodegeneration (65). AINX has no reported direct relationship to ALS pathology beyond its identity as an intermediate filament and its ability to co-assemble with NFL and NFM in mature CNS (41,44,45).

**Table 2:** The top features in the raw and BDC-ComBat-corrected datasets, as determined by principal component analysis. Protein and gene names are included. The number in parenthesis indicates whether the protein was a top feature for PC 1 or PC 2. The last column describes the general function of each feature.

| Protein/Gene | PCA (Raw) | PCA (BDC-ComBat) | Function |
|---|---|---|---|
| **AINX (INA)** | 0.854 (1) | 0.889 (1) | Alpha-internexin: Class-IV neuronal intermediate filament that can self-assemble, involved in the morphogenesis of neurons, may work with NFL/NFM/NFH |
| **NFL (NEFL)** | 0.263 (1) | 0.189 (1) | Neurofilament light polypeptide: Neurofilaments usually contain three intermediate filament proteins which are involved in the maintenance of neuronal caliber |
| **NFM (NEFM)** | 0.252 (1) | 0.178 (1) | Neurofilament medium polypeptide: Neurofilaments usually contain three intermediate filament proteins which are involved in the maintenance of neuronal caliber |
| **ENOA (ENO1)** | 0.359 (2) | NA | Alpha-enolase: Glycolytic enzyme the catalyzes the conversion of 2-phosphoglycerate to phosphoenolpyruvate |
| **QOR (CRYZ)** | NA | 0.220 (2) | Quinone oxidoreductase/Crystallin zeta: Crystallin protein which has NADPH-dependent quinone reductase activity distinct from other known quinone reductases |
| **KCRB (CK)** | 0.350 (2) | NA | Creatine kinase B-type: Reversibly catalyzes the transfer of phosphate between ATP and various phosphogens |
| **H15 (H1-5)** | 0.273 (2) | 0.853 (2) | Histone H1.5: Histones are basic nuclear proteins that are responsible for the nucleosome structure of the chromosomal fiber in eukaryotes |

The top-contributing protein to PC2 in the BDC-ComBat-corrected dataset is H15 (Histone 1.5) which has not been demonstrated as related to ALS. The second-most contributing protein in PC2 is AINX, like PC1. The third-most contributing protein to PC2 is quinone oxidoreductase (QOR), otherwise known as crystallin zeta (CRYZ) (42). There is little evidence to link QOR to ALS, and it is not specifically nor highly expressed in the central nervous system (43,44). However, one study that did a two-step genome-wide association study (GWAS) in Chinese Han people with ALS and controls did find that the TYW/CRYZ gene locus was associated with ALS (66). More research is needed to support a clear relationship between QOR/CRYZ and ALS.

## Discussion

The Answer ALS Consortium began collecting data from participants in 2016 and the first proteomes were released in 2021 (36). The iMDBs were generated in chronological order, meaning that iMDB 10 was differentiated earlier in time, and iMDB 60 was differentiated most recently (T. Thompson, personal communication, March 30^th^ 2026). While the cause(s) of the batch effect are unknown, changes in sample collection, iPSC differentiation, proteomic sequencing, or computer platform processing could have occurred in the collection time frame and contributed to a batch effect. Regardless of the cause, batch effect is not an uncommon problem. Many projects with longer study periods have been found to contain some kind of batch effect (28–31). Our findings provide support for workable correction methods that can help identify and mitigate batch effect in other large datasets.

Unique to the Answer ALS proteomic dataset was the outlier status of MOS and H4. Expression levels of these proteins correlated with PC1 and 2 when a PCA plot was generated from the raw proteomic dataset. The iMDB/shipment IDs of samples were also correlated with MOS and H4 expression, which indicates that MOS and H4 expression levels either correlate with batch effect or cause it. These proteins have not been previously reported as extreme outliers or major contributors to batch effect. MOS is a serine/threonine kinase involved in MAPK signaling, and H4 is Histone 4, a core component of the nucleosome (41). Neither protein is considered related to ALS, and neither is specifically nor highly expressed in the central nervous system (43,44). It is unclear why expressions of these two proteins would have such a large impact on the shape of the full proteomic dataset. However, we determined that MOS and H4 expression levels were not the sole cause of batch effect as removing both proteins from the dataset did not universally reduce batch effect. Further study is needed to determine why and how MOS and H4 expression levels are related to batch effect in the Answer ALS proteomic dataset, but it is our recommendation to remove them as outliers when analyzing this dataset.

We also determined that correcting the dataset with the internal non-disease control samples (BDCs) was insufficient to correct for batch effect. Samples in iMDBs 47-60 were artificially inflated when corrected with BDCs due to the BDCs being lower in average protein expression than any other samples in those iMDBs. Since iMDBs 10-45 were well-corrected by BDCs alone, it is possible to simply remove the later iMDBs and proceed without doing further correction. However, removing the samples in iMDBs 47-60 reduces the already-reduced dataset from 406 proteomic samples to 309 samples. It is also important to consider the future of the Answer ALS proteomic dataset, as iMDBs are planned for future release up to iMDB 99. Any correction method needs to be applicable to the current dataset as well as the future version of the dataset, of which the proteomic samples are still uncharacterized.

Towards our efforts to find a suitable correction method, we found that BDC-ComBat correction is more effective at reducing batch effect, both globally and in individual proteins, than BDC correction alone or ComBat-BDC correction. Additionally, BDC-ComBat correction preserved more of the original dataset than the other two approaches. No significant differences in protein expression levels between ALS case samples and control samples were found in any of the top PC-contributing proteins when BDC-ComBat correction was applied. This is an expected finding, considering that we determined batch effect is a major contributor to the variability of the dataset and then removed the batch effect. The lack of differences between cases and controls is also a finding reported by Workman et al., who were unable to meaningfully separate cases from controls in the Answer ALS transcriptomic dataset (25).

Four out of six features that contribute highly to PCs 1 and 2 in the BDC-ComBat-corrected dataset are at least tangentially related to ALS pathology, and three are highly and specifically expressed in brain tissues. This suggests that the batch effect has been reduced enough to discern genuine biology within the proteomic results. Additionally, the BDC-ComBat-corrected dataset recapitulates four of the six features that contribute highly to PCs 1 and 2 in the uncorrected dataset. This suggests that while the BDC-ComBat correction method reduces batch effect, the underlying structure of the proteomic dataset is preserved.

## Conclusion

In this study, we confirmed the presence of a batch effect in the Answer ALS proteomic Dataset 1-2. The overall average protein expression and individual protein expressions in a sample correlated to the iMDB in which the sample was differentiated from an iPSC to a motor neuron. We tried to correct for batch effect using the batch differential controls (BDCs) provided by Answer ALS and were ultimately not successful in reducing the effect. Correcting with the BDCs and then the batch effect correction algorithm (BECA) ComBat was successful in reducing batch effect both globally and on individual protein features.

While we cannot guarantee that the batch effect is completely eliminated using this method, BDC-ComBat correction is the method that we recommend for future analysis of proteomes. The top PC-contributing proteins discovered using this method were possibly related to ALS and highly expressed in the central nervous system. This method best reduces batch effect globally and for individual proteins, requires the removal of the least number of samples, and preserves the top-contributing features of the original proteomic dataset.

The Answer ALS project is an incredibly rich resource. The use of proteomic data from Answer ALS is limited due to a discovered batch effect that impacts the interpretability of the dataset. Here, we provide a road map for navigating the proteomic Answer ALS dataset that can facilitate further exploration of this resource to advance research for this complex neurodegenerative disorder.

## Declarations

### Ethics approval and consent to participate

Data used in the preparation of this article were obtained from the ANSWER ALS Data Portal (AALS-01184). For up-to-date information on the study, visit https://dataportal.answerals.org. Access to the data was granted to the authors of this article through a Data Use Agreement (DUA) with Answer ALS signed in 2022. All Answer ALS participants were de-identified prior to our procurement of the data and remain de-identified in this article. When we refer to an individual participant, we use the Answer ALS generated ‘Participant ID’.

### Consent for publication

Not Applicable

### Availability of data and materials

Data used in the preparation of this article were obtained from the ANSWER ALS Data Portal (AALS-01184). For up-to-date information on the study, visit https://dataportal.answerals.org. The data that support the findings of this study are available from Answer ALS but restrictions apply to the availability of these data, which were used under license for the current study, and so are not publicly available. Data are however available from the authors upon reasonable request and with permission of Answer ALS. One exception is the ‘Sample Mapping File’ which describes the order in which samples were processed, and is publicly available here, https://dataportal.answerals.org/data-information, under the tab ‘Sample and Batch Mapping Information’. This file is also available in our Supplemental Materials.

### Authors’ contributions

GC performed all experiments in this study, created all figures and tables, and wrote the majority of this manuscript. IO helped analyze research findings and determine the appropriate experimental outline. ZA helped analyze experimental findings and determine the appropriate experimental outline. PP helped obtain access to the Answer ALS data and provided mentorship. TMM helped obtain access to the Answer ALS data and provided mentorship. All authors contributed to the editing of this manuscript, and all authors read and approved the final manuscript.

## Acknowledgements

We thank the participants in Answer ALS who contributed to this dataset and also participated in multiple interviews and clinical tests to provide as much information as possible, all while having a disorder that severely limits mobility. We thank the Program Director of OnPoint Scientific, Dr. Terri Thompson, for answering our questions about the Answer ALS data generation process. We also thank Dr. Kathleen Schoch for editing many iterations of this manuscript.

