## Supplemental Figures 1-17 for "Batch Effect Correction to Enhance Utility of a Publicly Available ALS Proteomics Dataset"

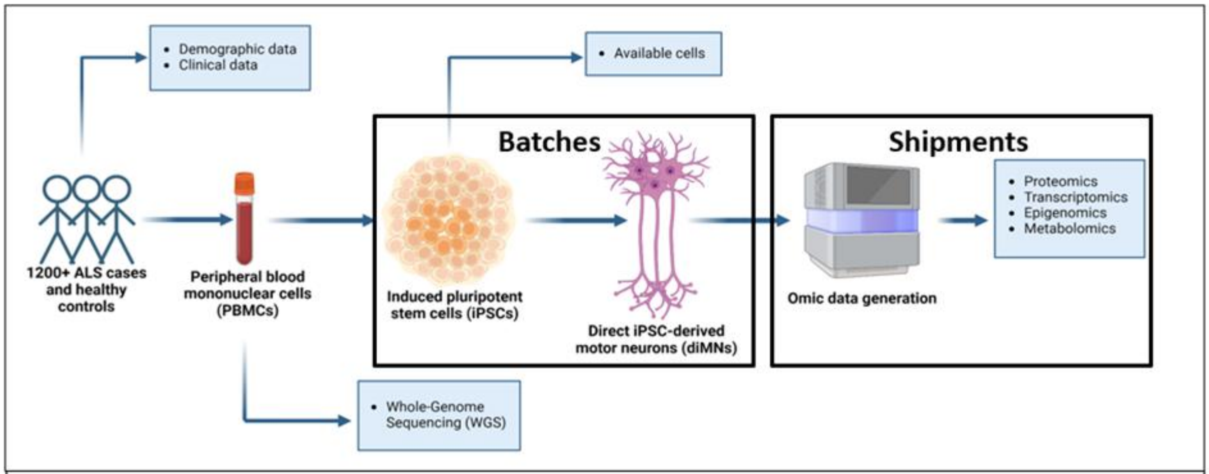


**Supplemental Figure 1: The process of proteomic data generation by Answer ALS.** Participant sample peripheral blood mononuclear cells (PBMCs) were differentiated into induced pluripotent stem cells (iPSCs), then differentiated again into direct iPSC-derived motor neurons (diMNs), in batches of 10-13 samples. Mass spectrometry was then performed at Cedars-Sinai Medical Center on 2-4 batches of diMNs at one time, referred to as a 'shipment'.

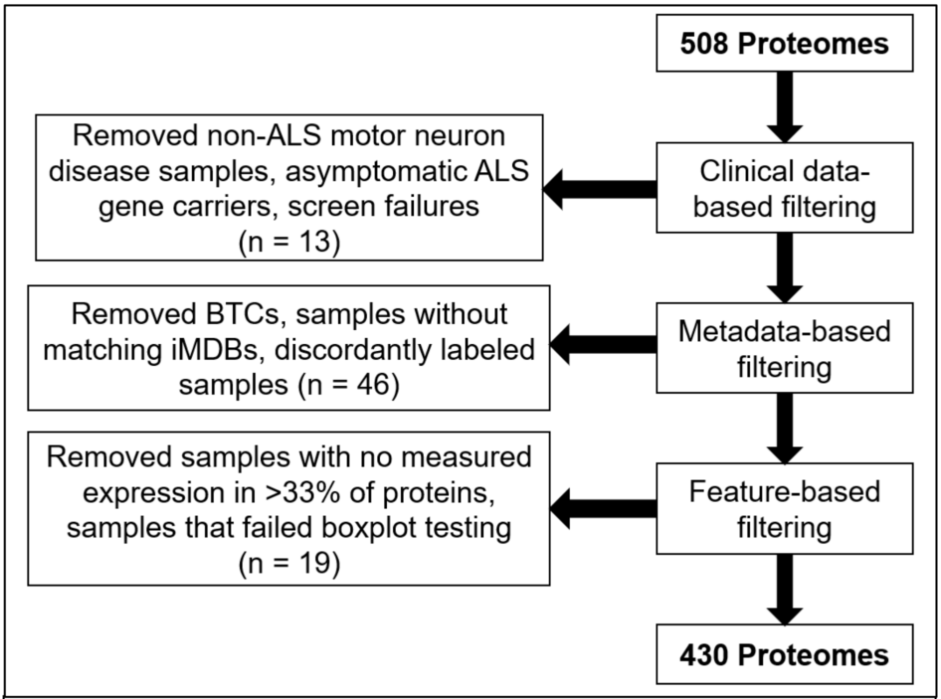


**Supplemental Figure 2: Filters applied to the Answer ALS proteomic dataset.** 50 proteomic samples were removed through clinical data, metadata, and feature-based filters.

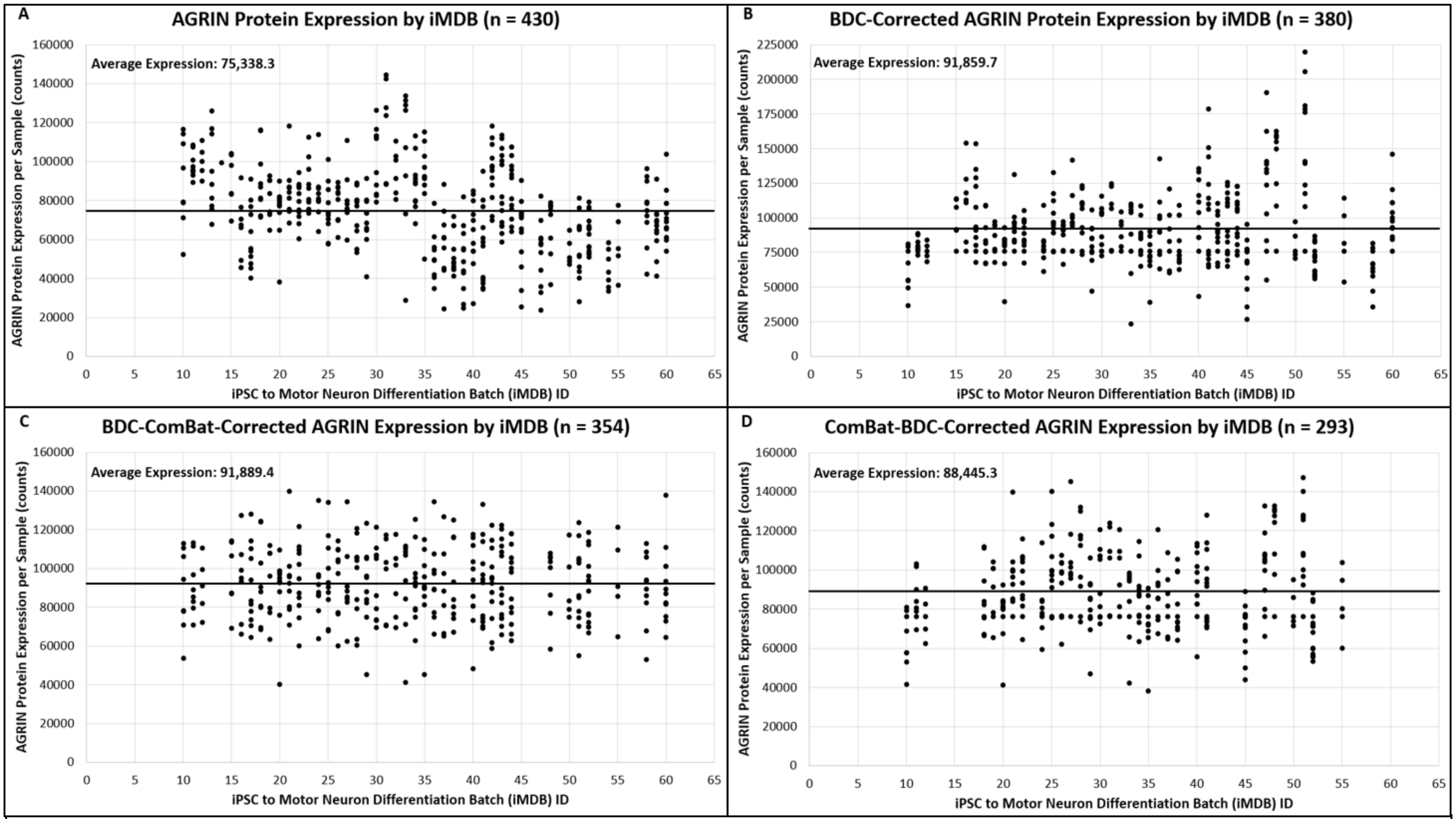


**Supplemental Figure 3: The distribution of random protein 1, AGRIN, across iMDBs in four different iterations of the proteomic dataset.** AGRIN protein expression per sample across iMDBs in the A) raw proteomic dataset, B) BDC-corrected dataset, C) BDC-ComBat-corrected dataset, and D) the ComBat-BDC-corrected dataset.

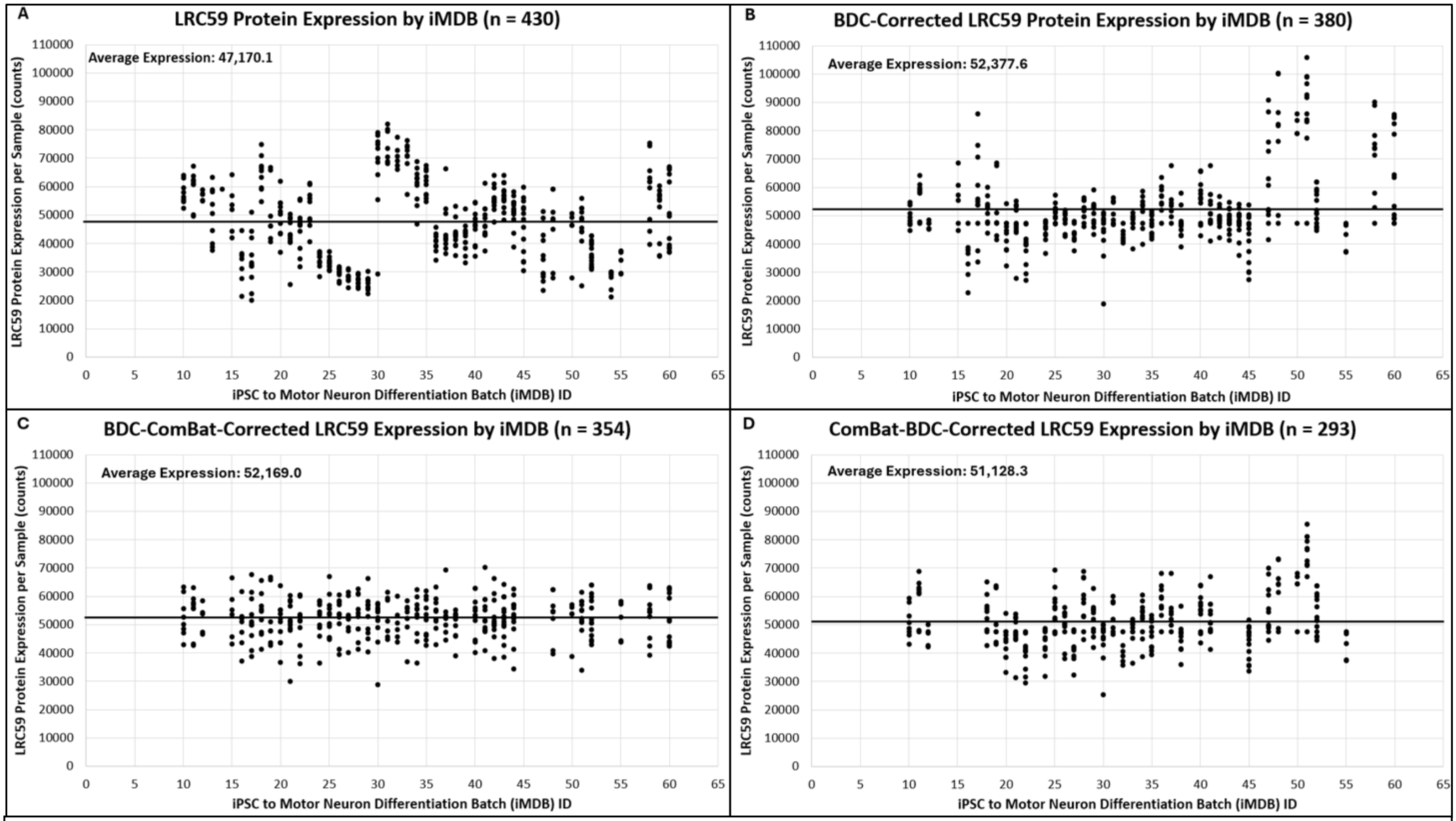


**Supplemental Figure 4: The distribution of random protein 2, LRC59, across iMDBs in four different iterations of the proteomic dataset.** LRC59 protein expression per sample across iMDBs in the A) raw proteomic dataset, B) BDC-corrected dataset, C) BDC-ComBat-corrected dataset, and D) the ComBat-BDC-corrected dataset.

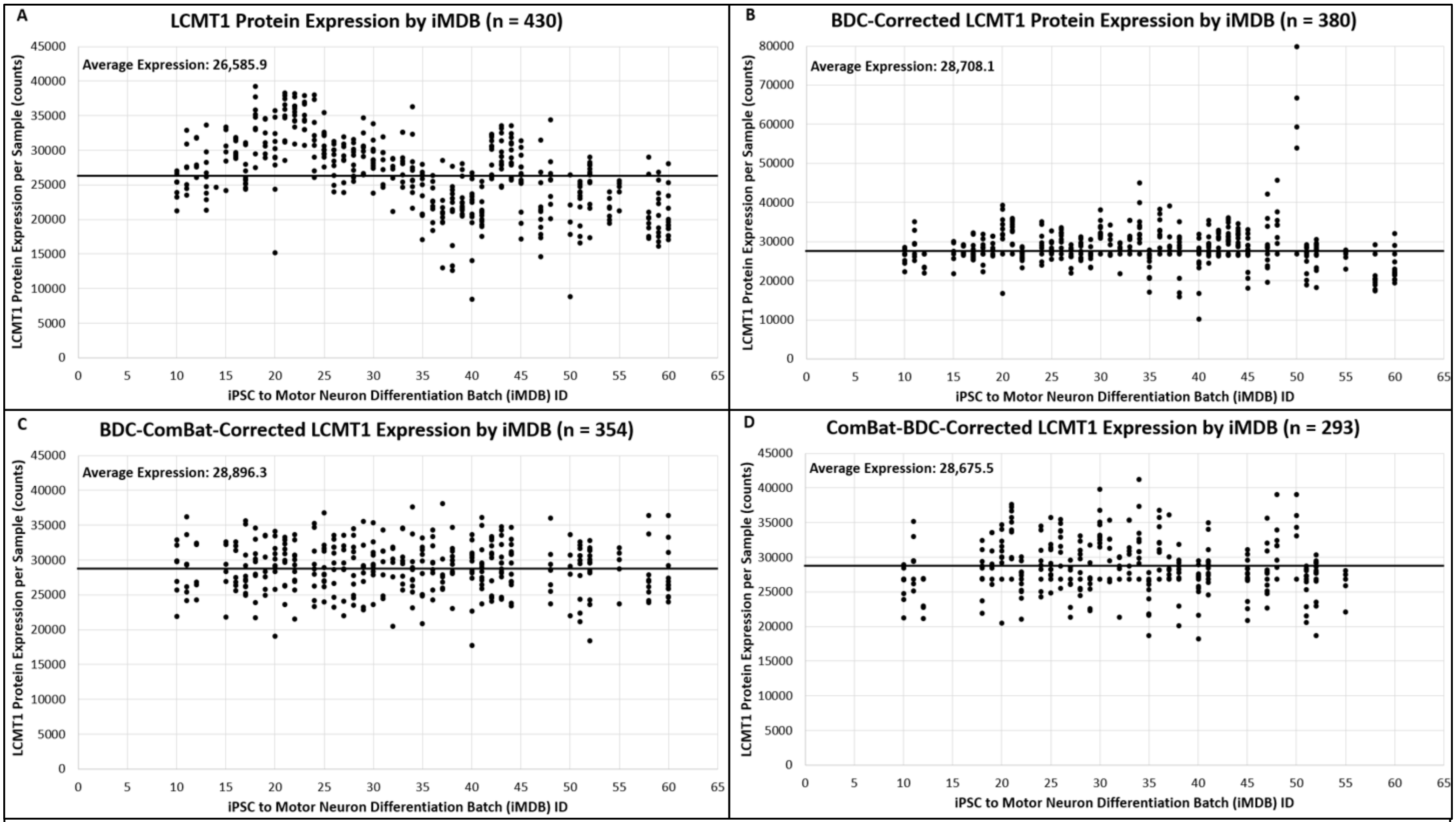


**Supplemental Figure 5: The distribution of random protein 3, LCMT1, across iMDBs in four different iterations of the proteomic dataset.** LCMT1 protein expression per sample across iMDBs in the A) raw proteomic dataset, B) BDC-corrected dataset, C) BDC-ComBat-corrected dataset, and D) the ComBat-BDC-corrected dataset.

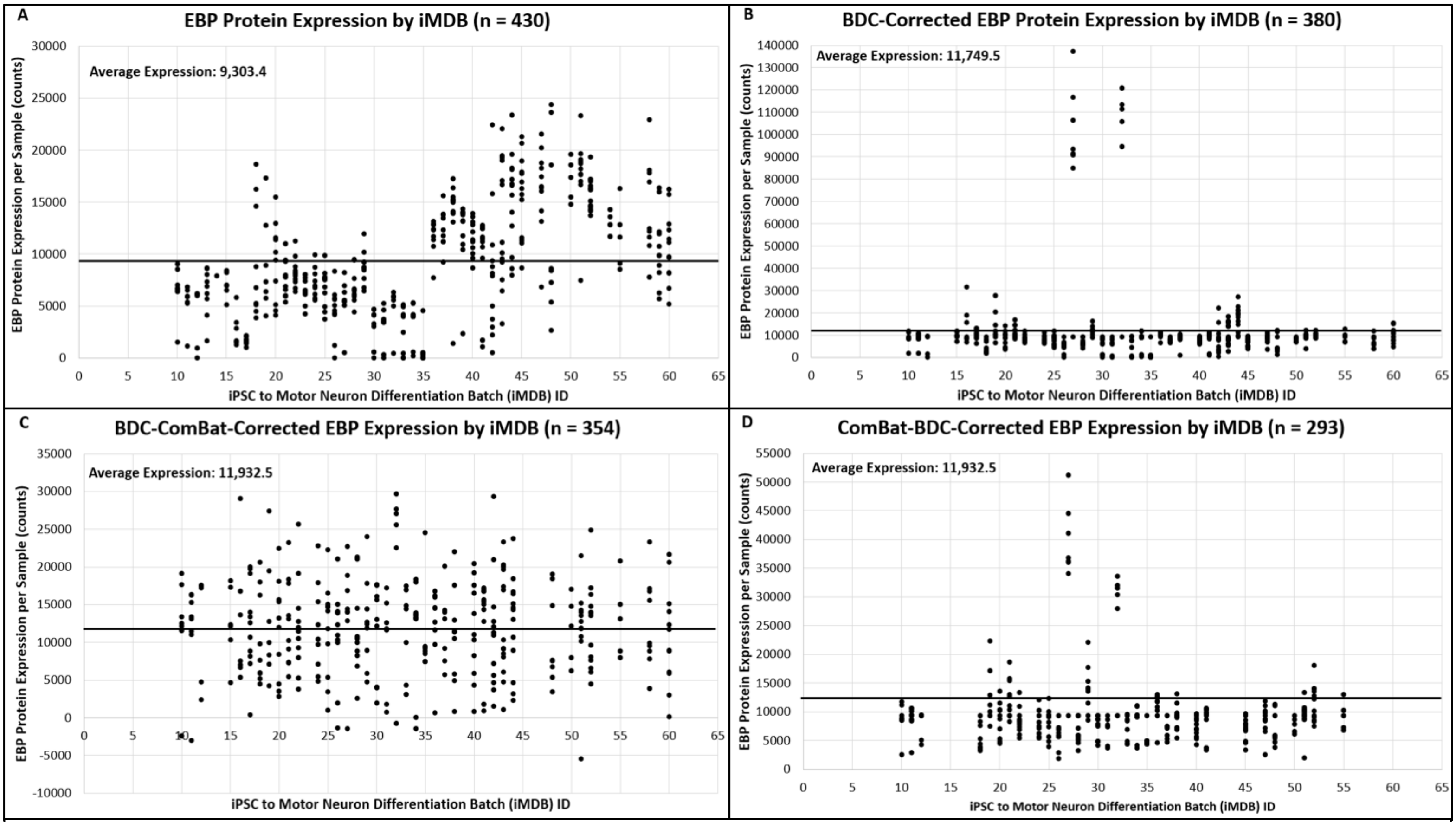


**Supplemental Figure 6: The distribution of random protein 4, EBP, across iMDBs in four different iterations of the proteomic dataset.** EBP protein expression per sample across iMDBs in the A) raw proteomic dataset, B) BDC-corrected dataset, C) BDC-ComBat-corrected dataset, and D) the ComBat-BDC-corrected dataset.

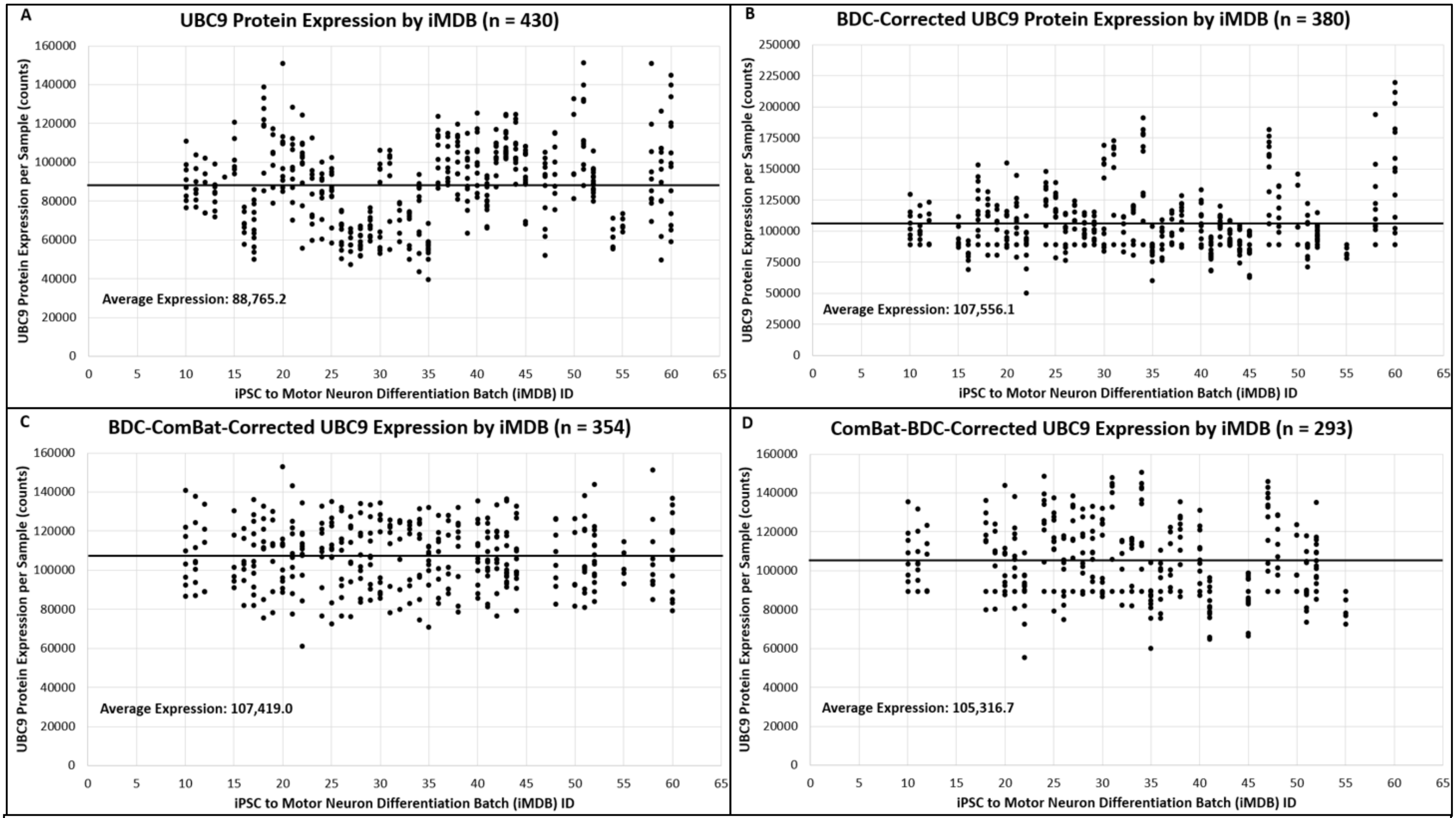


**Supplemental Figure 7: The distribution of random protein 5, UBC9, across iMDBs in four different iterations of the proteomic dataset.** UCB9 protein expression per sample across iMDBs in the A) raw proteomic dataset, B) BDC-corrected dataset, C) BDC-ComBat-corrected dataset, and D) the ComBat-BDC-corrected dataset.

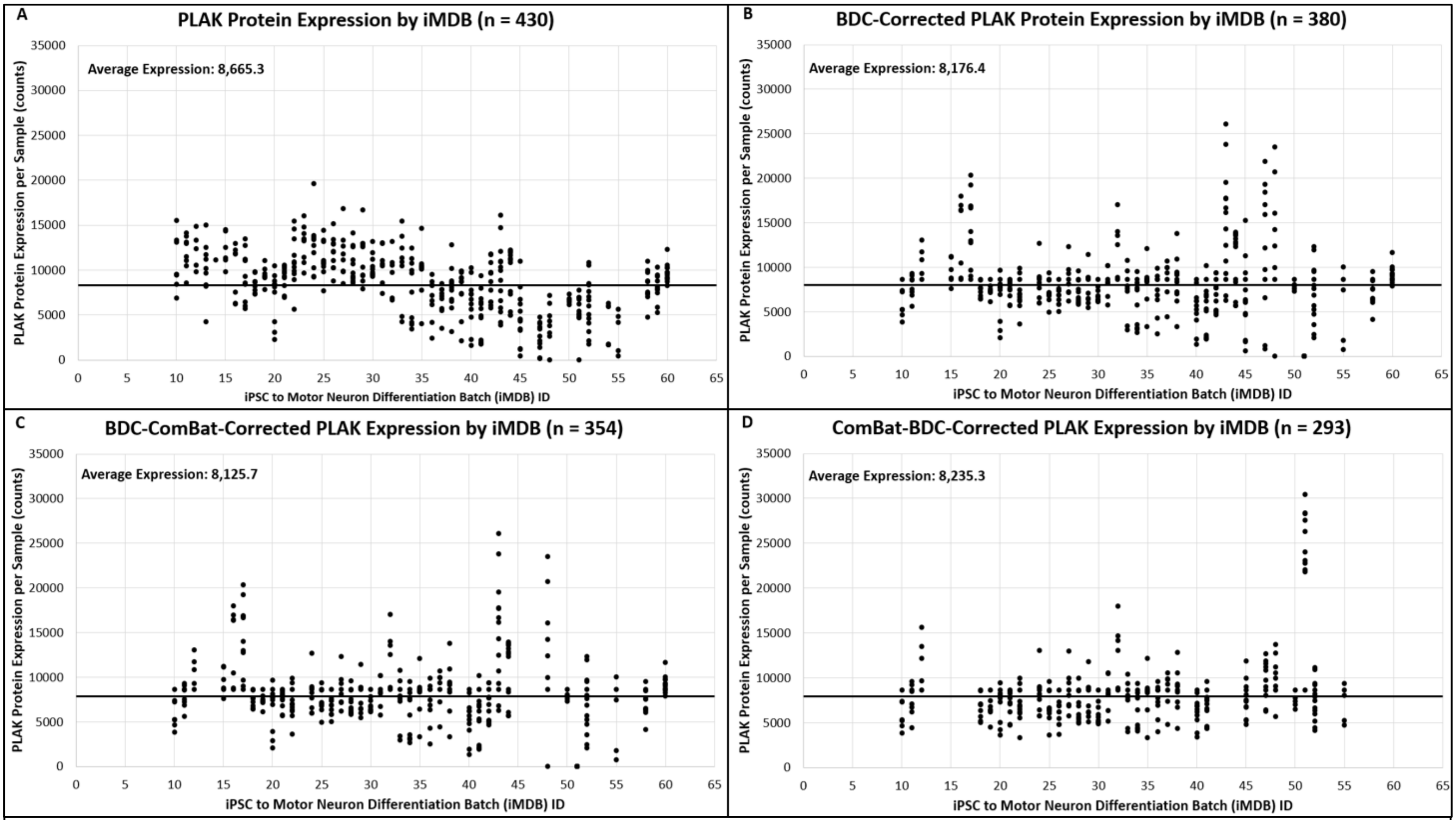


**Supplemental Figure 8: The distribution of random protein 6, PLAK, across iMDBs in four different iterations of the proteomic dataset.** PLAK protein expression per sample across iMDBs in the A) raw proteomic dataset, B) BDC-corrected dataset, C) BDC-ComBat-corrected dataset, and D) the ComBat-BDC-corrected dataset.

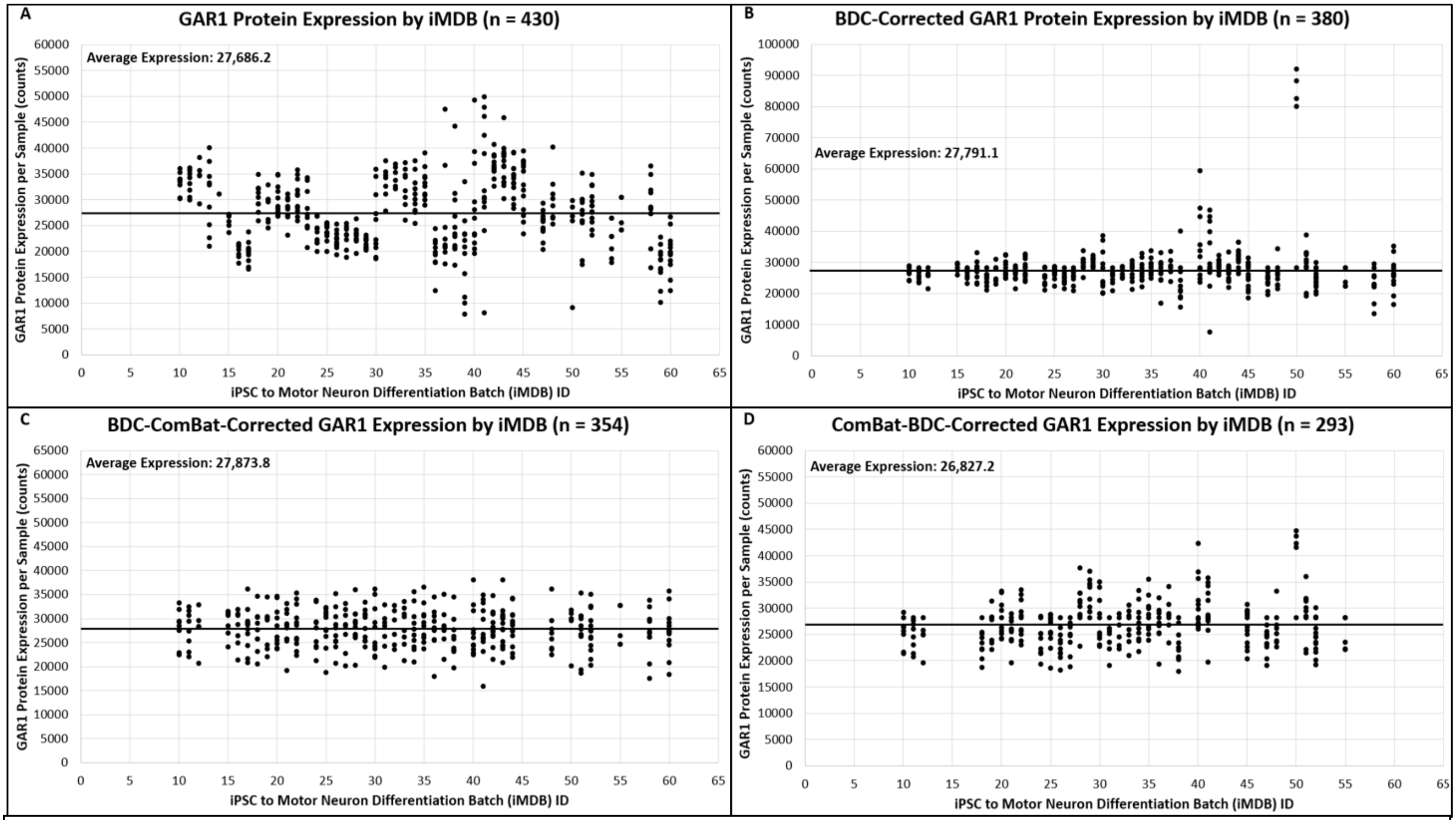


**Supplemental Figure 9: The distribution of random protein 7, GAR1, across iMDBs in four different iterations of the proteomic dataset.** GAR1 protein expression per sample across iMDBs in the A) raw proteomic dataset, B) BDC-corrected dataset, C) BDC-ComBat-corrected dataset, and D) the ComBat-BDC-corrected dataset.

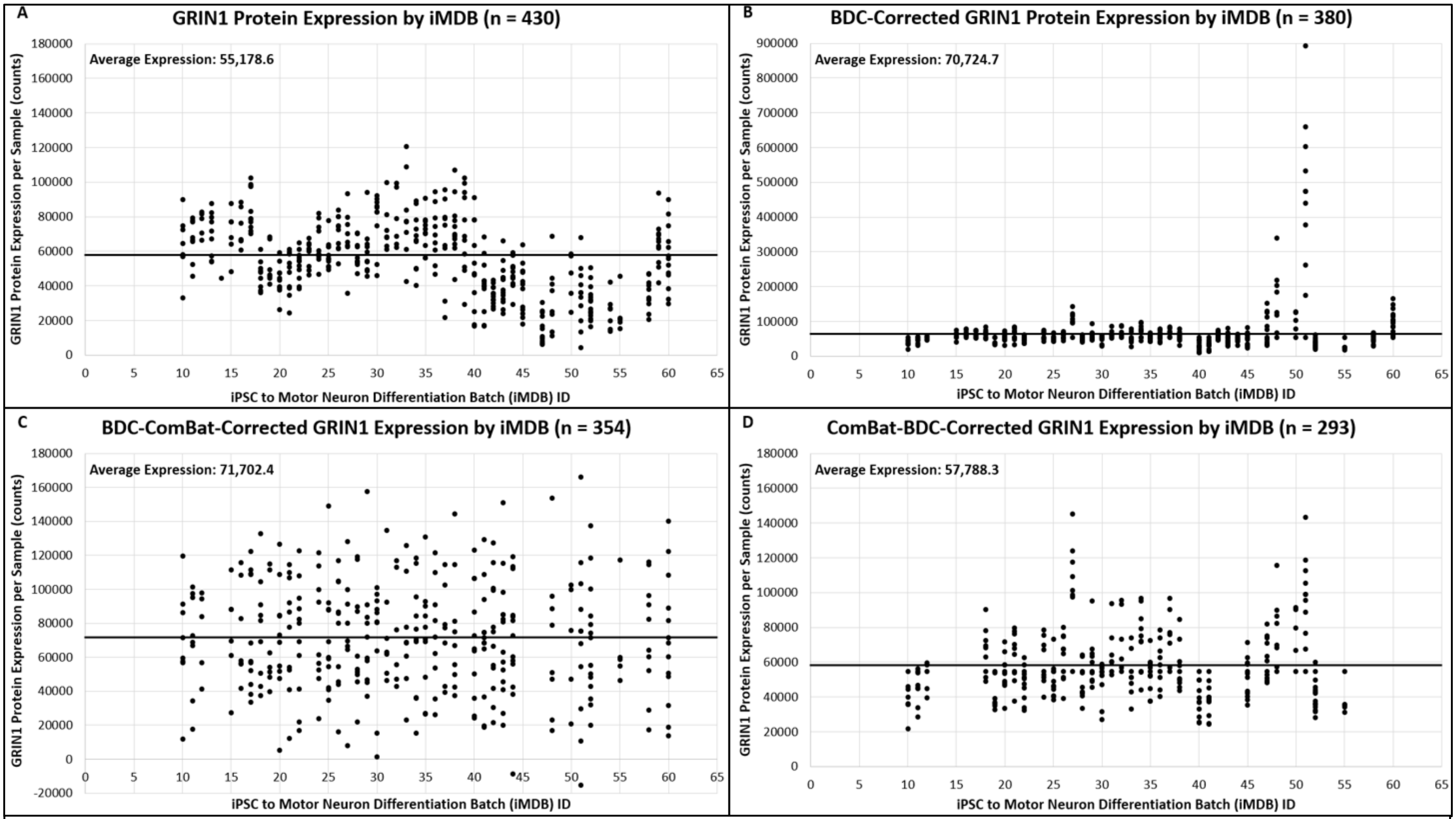


**Supplemental Figure 10: The distribution of random protein 8, GRIN1, across iMDBs in four different iterations of the proteomic dataset.** GRIN1 protein expression per sample across iMDBs in the A) raw proteomic dataset, B) BDC-corrected dataset, C) BDC-ComBat-corrected dataset, and D) the ComBat-BDC-corrected dataset.

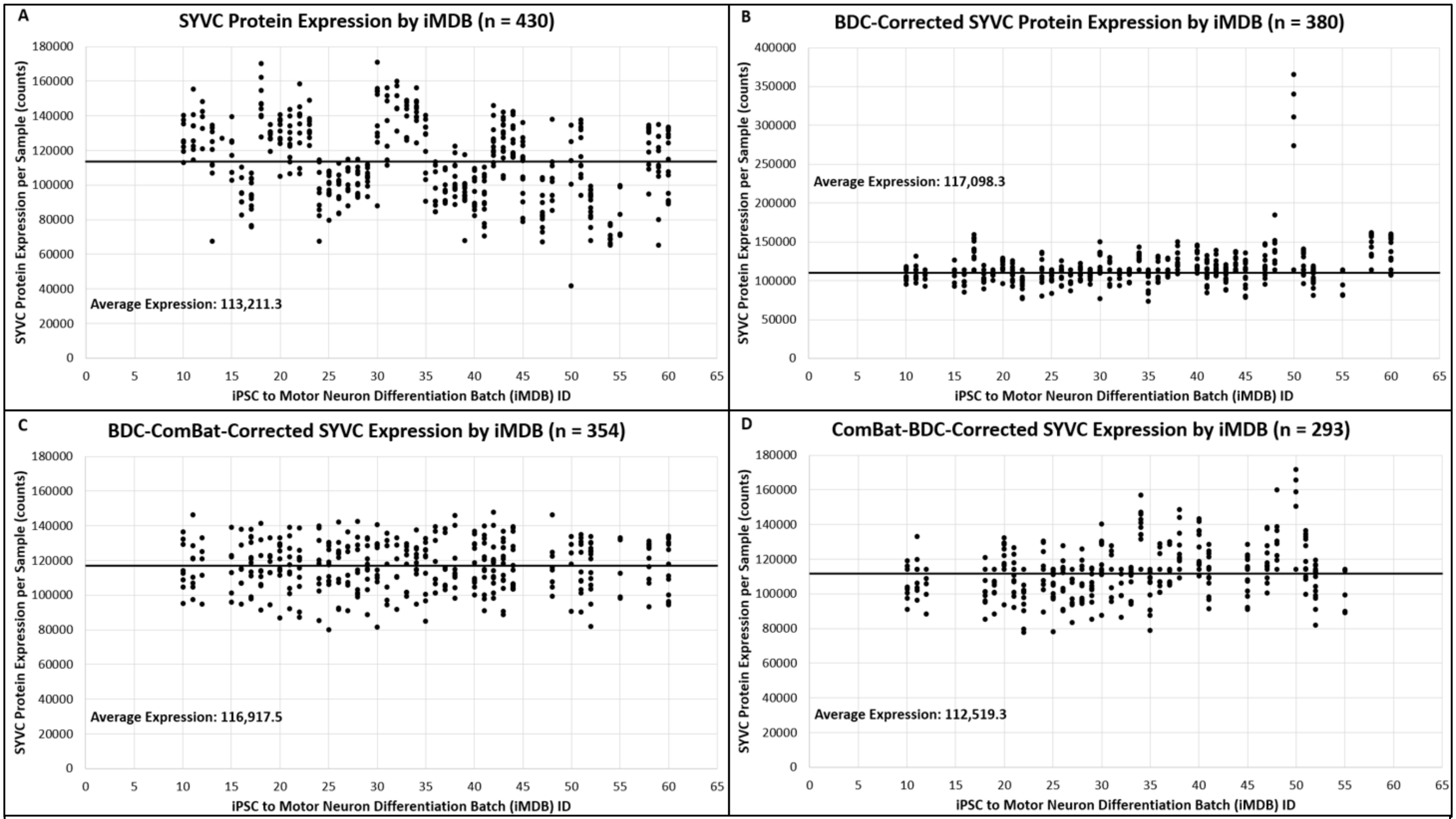


**Supplemental Figure 11: The distribution of random protein 9, SYVC, across iMDBs in four different iterations of the proteomic dataset.** SYVC protein expression per sample across iMDBs in the A) raw proteomic dataset, B) BDC-corrected dataset, C) BDC-ComBat-corrected dataset, and D) the ComBat-BDC-corrected dataset.

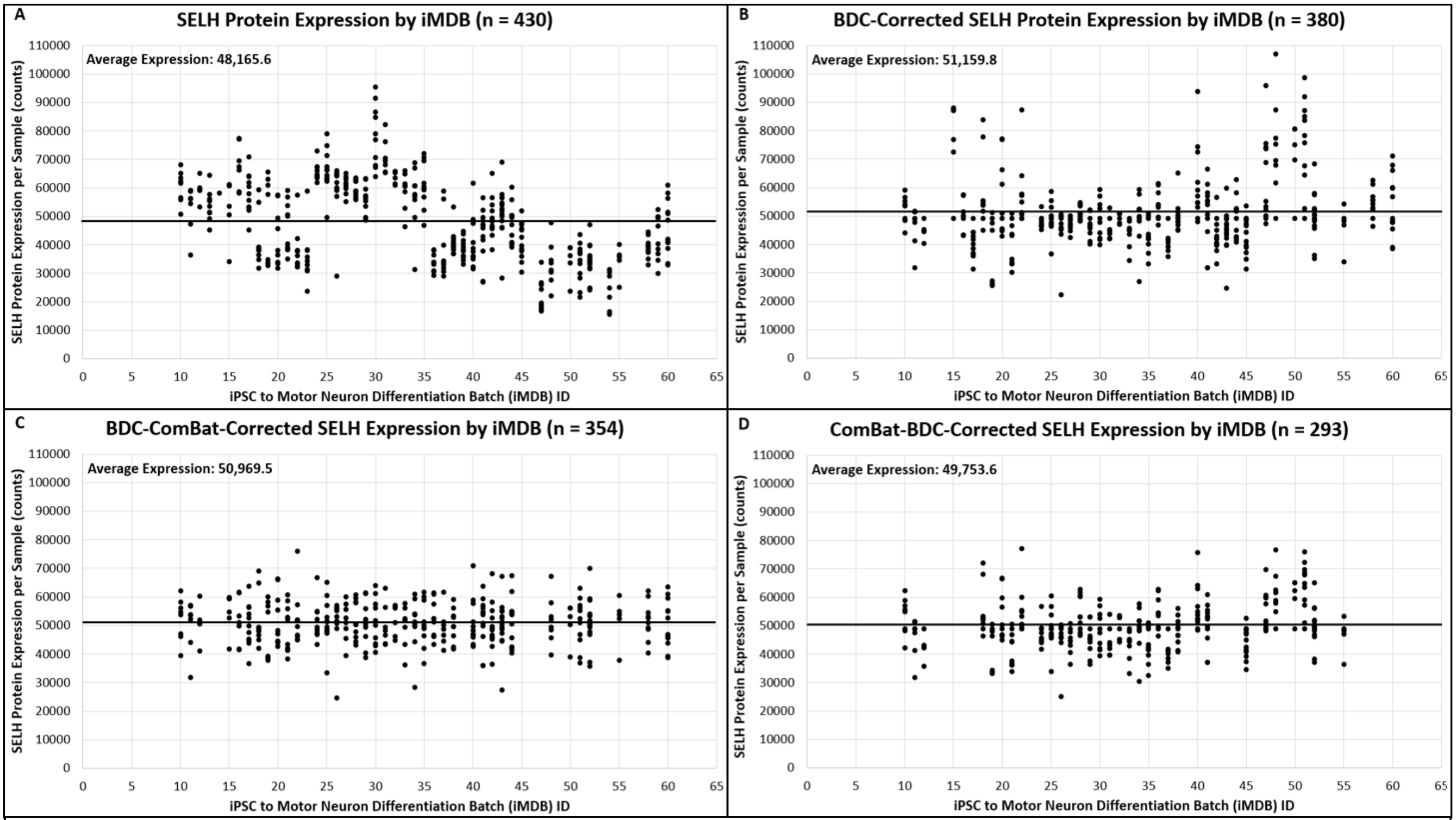


**Supplemental Figure 12: The distribution of random protein 10, SELH, across iMDBs in four different iterations of the proteomic dataset.** SELH protein expression per sample across iMDBs in the A) raw proteomic dataset, B) BDC-corrected dataset, C) BDC-ComBat-corrected dataset, and D) the ComBat-BDC-corrected dataset.

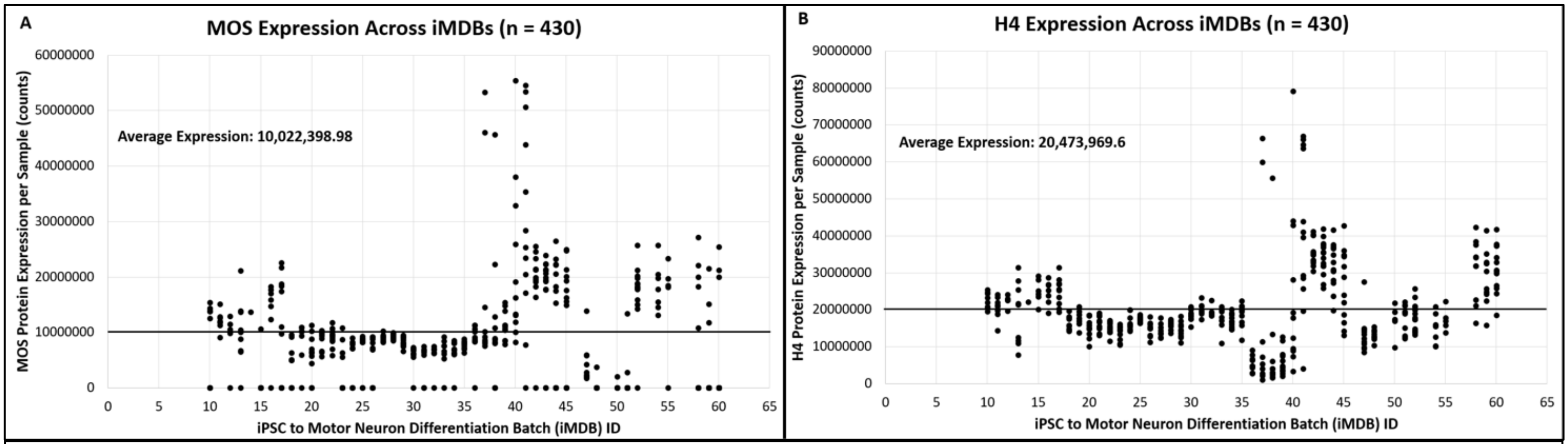


**Supplemental Figure 13: The distribution of outlier proteins MOS and H4 across iMDBs in the original proteomic dataset.** A) MOS protein expression per sample across iMDBs and B) H4 protein expression per sample across iMDBs.

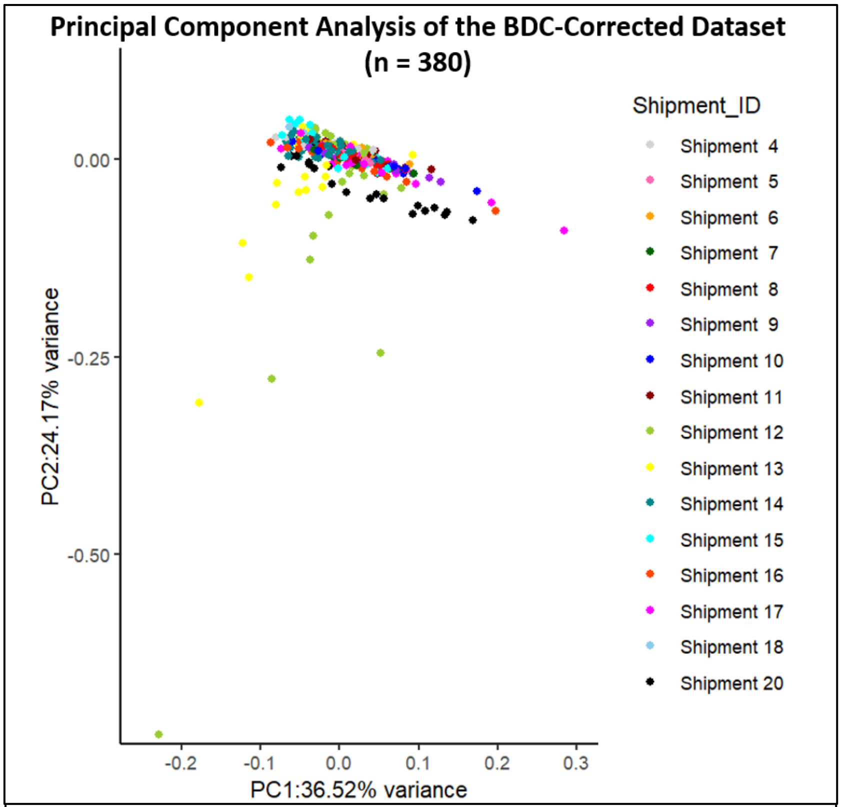


**Supplemental Figure 14: An additional outlier sample was identified after performing PCA with the other outliers removed.** PCA plot of 380 BDC-corrected samples where 'NEUWD946RDA' was identified as an outlier, color-coded by shipment ID.

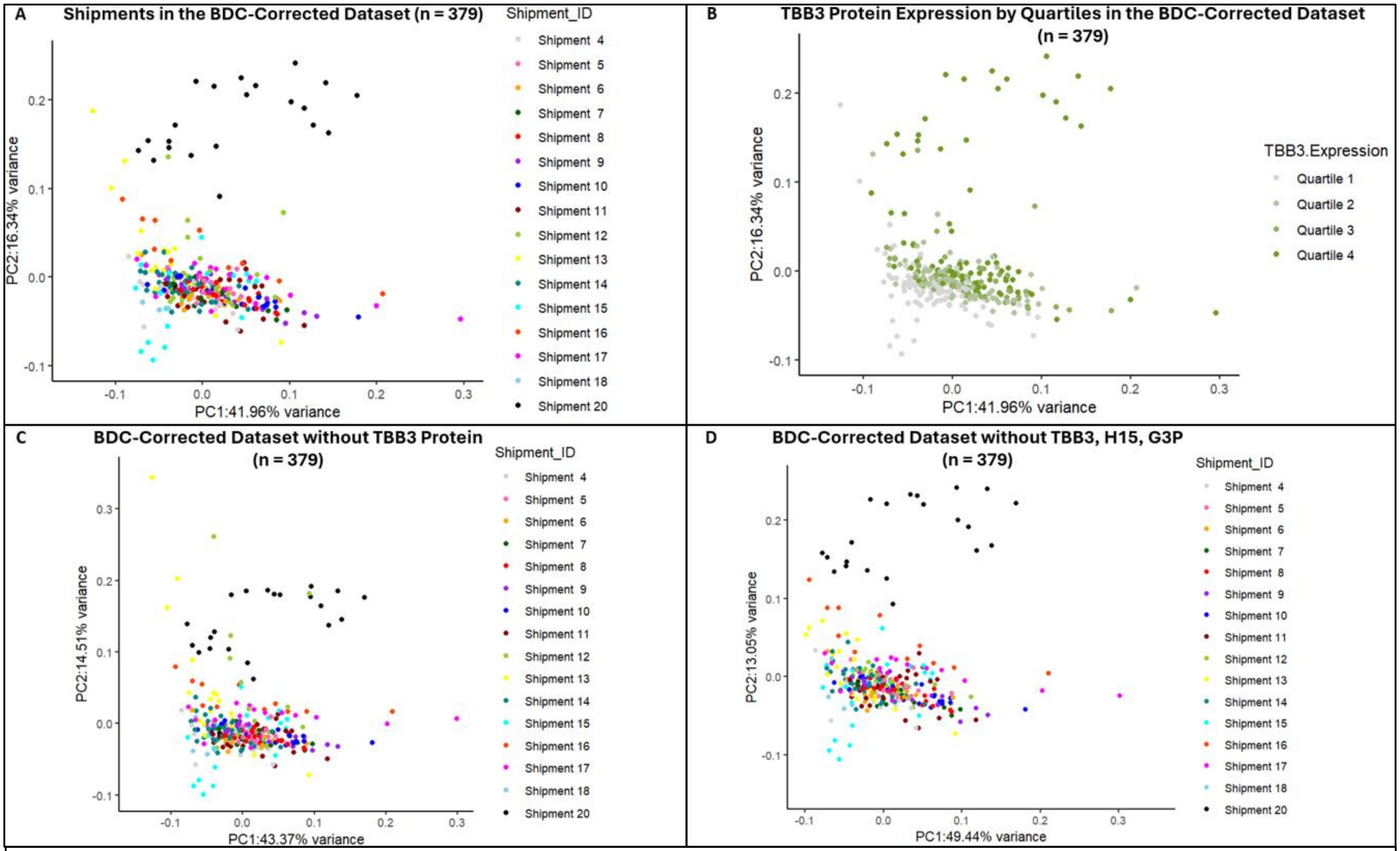


**Supplemental Figure 15: Removing top PC-contributing proteins in the BDC-corrected dataset is not sufficient for correcting batch effect.** A) PCA plot of BDC-corrected proteomic dataset (n = 3,641) with outlier samples removed, color-coded by shipment ID. B) PCA plot of the BDC-corrected dataset color-coded by quartile expression of TBB3, the highest contributing protein to PC2. C) PCA plot of the BDC-corrected dataset with TBB3 removed from the dataset (n = 3,640), color-coded by shipment ID. D) PCA plot of the BDC-corrected dataset with TBB3, H15, and G3P (the top three highest contributing protein to PC2) removed from the dataset (n = 3,638), color-coded by shipment ID.

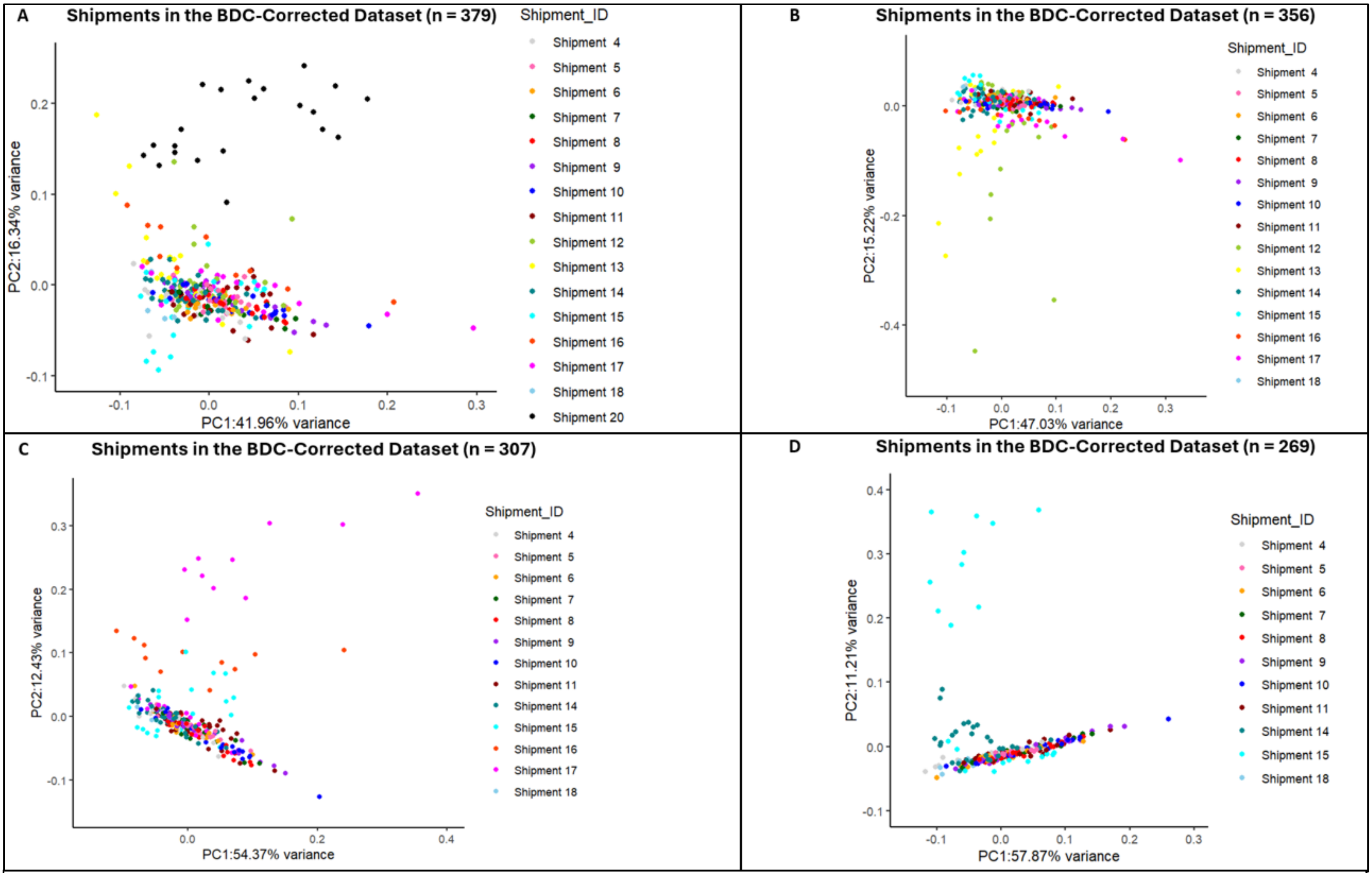


**Supplemental Figure 16: Removing outlier shipments/iMDBs in the BDC-corrected dataset is not sufficient for correcting batch effect.** A) PCA plot of BDC-corrected proteomic dataset color-coded by shipment ID. B) PCA plot of the BDC-corrected dataset with shipment 20 (iMDBs 58-60) removed, color-coded by shipment ID. C) PCA plot of the BDC-corrected dataset with shipments 12, 13, and 20 (iMDBs 36-38, 40, 41, 58-60) removed, color-coded by shipment ID. D) PCA plot of the BDC-corrected dataset with shipments 12, 13, 16, 17, and 20 (iMDBs 36-38, 40, 41, 48, 50-52, 58-60) removed, color-coded by shipment ID.

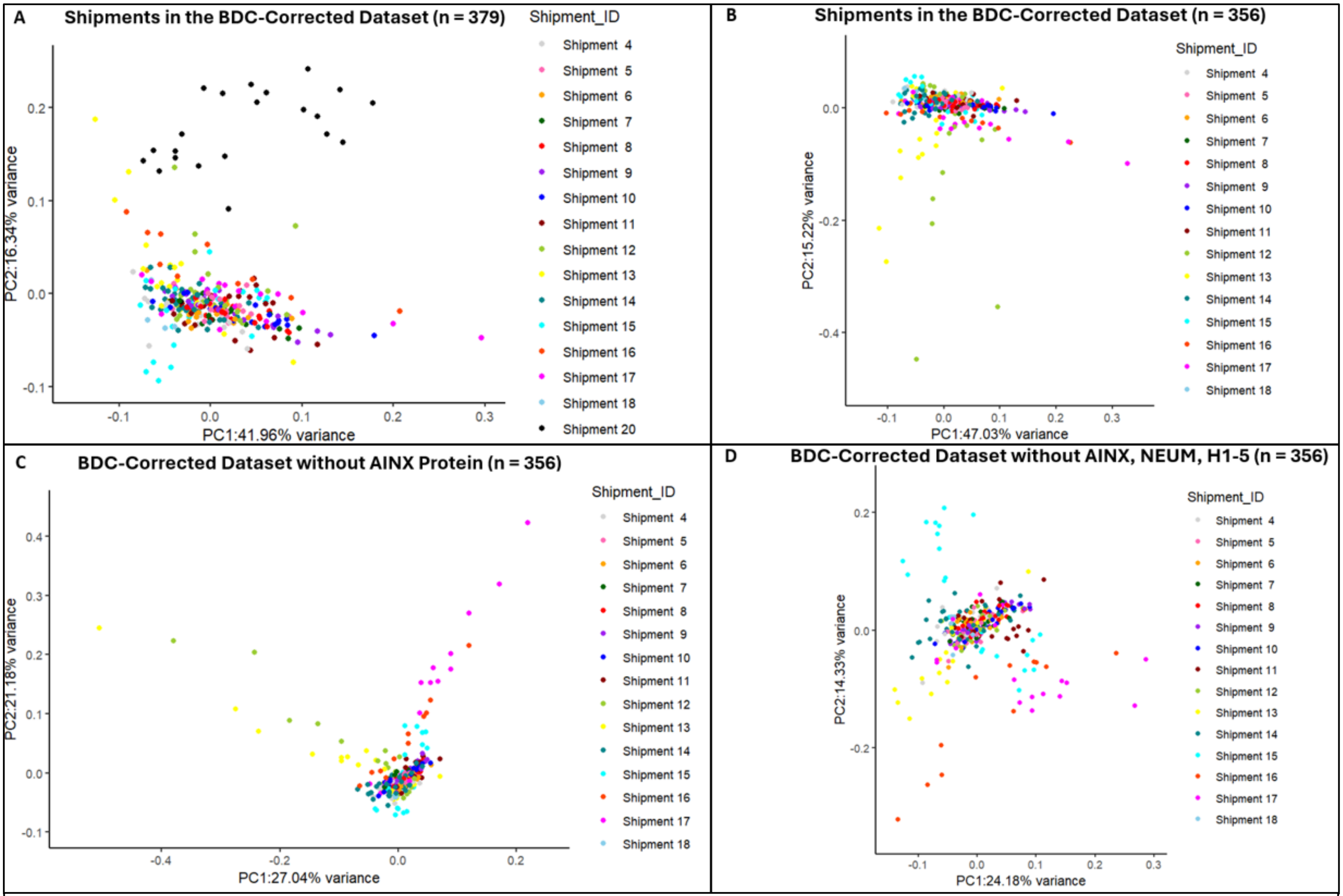


**Supplemental Figure 17: Removing outlier shipments/iMDBs and top PC-contributing proteins in combination in the BDC-corrected dataset is not sufficient for correcting batch effect.** A) PCA plot of BDC-corrected proteomic dataset color-coded by shipment ID. B) PCA plot of the BDC-corrected dataset with shipment 20 (iMDBs 58-60) removed, color-coded by shipment ID. C) PCA plot of the BDC-corrected dataset with shipments 20 (iMDBs 58-60) and top PC2-contributing protein TBB3 removed from the dataset (n = 3,640), color-coded by shipment ID. D) PCA plot of the BDC-corrected dataset with shipment 20 (iMDBs 58-60) and the top three PC2-contributing proteins, TBB3, NEUM, and H15 removed from the dataset (n = 3,638), color-coded by shipment ID.
